# Fast searches of large collections of single cell data using scfind

**DOI:** 10.1101/788596

**Authors:** Jimmy Tsz Hang Lee, Nikolaos Patikas, Vladimir Yu Kiselev, Martin Hemberg

## Abstract

Single cell technologies have made it possible to profile millions of cells, but for these resources to be useful they must be easy to query and access. To facilitate interactive and intuitive access to single cell data we have developed scfind, a search engine for cell atlases. Using transcriptome data from mouse cell atlases we show how scfind can be used to evaluate marker genes, to perform *in silico* gating, and to identify both cell-type specific and housekeeping genes. Moreover, we have developed a subquery optimization routine to ensure that long and complex queries return meaningful results. To make scfind more user friendly and accessible, we use indices of PubMed abstracts and techniques from natural language processing to allow for arbitrary queries. Finally, we show how scfind can be used for multi-omics analyses by combining single-cell ATAC-seq data with transcriptome data.

## Introduction

Single cell technologies have made it possible to profile large numbers of cells (Cao et al., 2019; Cusanovich et al., 2018; Han et al., 2018; Saunders et al., 2018; Tabula Muris Consortium et al., 2018; Zeisel et al., 2018), and there are currently several ongoing efforts to build comprehensive atlases of humans (Regev et al., 2017) and other organisms (Cao et al., 2019; Han et al., 2018; Howick et al., 2019; Tabula Muris Consortium et al., 2018). One ambition of these projects is to create references that can be used as a foundation for both basic research and clinical applications (Regev et al., 2017). To achieve this goal and to ensure that cell atlases can fulfill their potential, it is critical that they are easy to access for many users, ranging from computational biologists interested in large scale analyses to wet lab biologists and clinicians who are interested in a specific gene, pathway, or disease. For example, a ChIP-seq experiment carried out on a tissue provides a list of genes targeted by a transcription factor, and the researcher may be interested in pinpointing the specific cell type that is using the transcription factor. Another example is the interpretation of genetic variants that have been associated with a disease. By identifying which cell types express the gene carrying the variant of interest, it is possible to gain important mechanistic insights (Calderon et al., 2017; Campbell et al., 2017; Kanai et al., 2018). To assist researchers analyzing complex datasets, it should be easy to interface with other widely used resources, e.g. the genome wide association study (GWAS) catalog (MacArthur et al., 2017), the gene ontology (GO) knowledgebase (The Gene Ontology Consortium, 2017), the disease ontology Medical Subject Headings (MeSH) (Sewell, 1964), and collections of clinically important genetic variants (Cariaso and Lennon, 2012; Landrum et al., 2016).

Many of the challenges in working with single cell data stem from its large size. Typically, a single cell RNA-seq (scRNA-seq) dataset is represented as a gene-by-cell expression matrix with ∼20,000 genes and may include millions of cells. For epigenetic data, e.g. scATAC-seq, the situation is often worse as the matrix can have many more rows, each representing a peak. Even though the matrix is sparse, working with such a dataset places high demands on computer hardware. Many operations are not only time consuming, but require advanced bioinformatics skills. To allow for both interactive and high-throughput queries of a large cell atlas, novel algorithms and data structures are required.

To ensure that large single cell datasets can be accessed by a wide range of users, the underlying software must (i) allow for complex queries, (ii) take full advantage of the single-cell resolution of the data, (iii) support different types of assays besides RNA-seq, (iv) be fast and easy to use, (v) be possible to run interactively through a web browser to access large repositories of data, (vi) be possible to run locally by a user interested in analyzing their own data. However, none of the computational tools available today for collections of scRNA-seq datasets, e.g. Panglao (Franzén et al., 2019), the UCSC Cell Browser (Haeussler et al., 2019), scRNASeqDB (Cao et al., 2017), SCPortalen (Abugessaisa et al., 2018), and the EBI Single Cell Expression Atlas (Athar et al., 2019), provide the required functionality and versatility.

Here, we present scfind, a search engine that makes single cell data accessible to a wide range of users by enabling sophisticated queries for large datasets through an interface which is both very fast and familiar to users from any background. The central operation carried out by scfind is to identify the set of cells that express a set of genes or peaks (i.e. the query) specified by the user. By themselves, such searches can be very powerful as they can identify the cell type that is most enriched for cells corresponding to the query within hundredths of a second. Furthermore, we demonstrate how the fast searches opens up new possibilities for global analyses of cell atlases, e.g. for identifying housekeeping genes and tissue-specific genes. By taking advantage of information from PubMed abstracts and natural language processing techniques, scfind allows a user to input any type of query and automatically translates it into gene list which is used for the search. Finally, we show how scfind can be applied to both scRNA-seq and scATAC-seq atlases together to identify putative cell type specific enhancers.

## Results

### Efficient compression allows for fast queries with single cell resolution

The input for building an scfind index is one or more non-negative matrices where rows represent genes, peaks, or another feature which are used as query terms and each column represents a cell. There are no assumptions regarding normalization, but it is beneficial if the matrix is sparse. To build a compact index to store the matrix, scfind uses the row names as keys, and compresses each row in two steps. The first step, which is lossless, stores the indices of the cells with non-zero values as binary strings using Elias-Fano coding (Vigna, 2013). In the second, lossy step, the approximate values are stored with a user-defined precision (Fig 1a). The combination of lossless and lossy strategies means that the most important information, whether or not a key is present in a cell, is perfectly retained, whereas, only an approximation of the value is stored to save space. Since the number of bits used per entry in the second step is decided by the user, it is possible to tune the trade-off between compression and information loss. If the cells have been assigned labels, e.g. using unsupervised clustering or projection methods, these are stored and used to group the cells returned from a query. Importantly, the index is modular, making it possible to add new datasets without having to reprocess the ones that have already been indexed.

**Figure 1:**
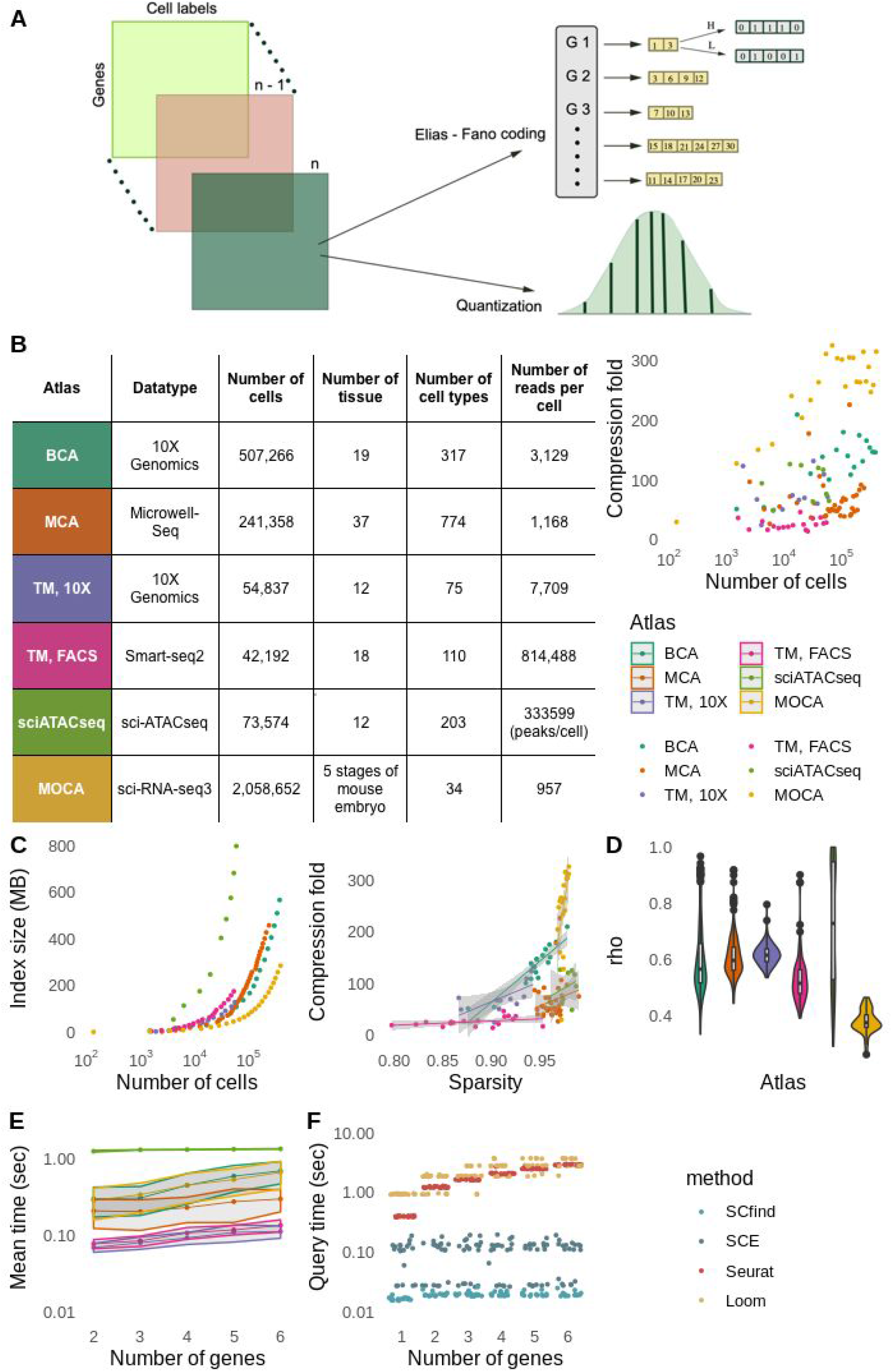
Compression and search times. (a) Schematic illustration of scfind’s compression strategy. Each square represents an expression matrix from an individual experiment. Each matrix is compressed and the resulting vectors are then concatenated. (b) Summary statistics for the six atlases considered in this manuscript along with compression ratios where each point represents one tissue. (c) Cumulative size of the resulting index when merging the different tissues (left) and fraction of zeros compared to compression ratios (right). For legend, see (b). (d) Accuracy of the reconstructed expression values when using two bits for the quantization. The violin plots show Spearman correlations for each gene and cell type. (e) Mean search times for AND queries involving up to six genes. Grey bands indicate standard errors for 100 repetitions. (f) Comparison of query times with three other file formats.

We applied scfind to five large mouse scRNA-seq datasets: the Mouse Organogenesis Cell Atlas (MOCA) (Cao et al., 2019), the Mouse Cell Atlas (MCA) (Han et al., 2018), two collections, FACS and 10X, from the Tabula Muris (TM) (The Tabula Muris Consortium et al., 2017) which are sampled from the whole body and the Zeisel (Zeisel et al., 2018) dataset, which contains cells from the brain. We also indexed the sciATAC-seq mouse atlas (Cusanovich et al., 2018) using the genomic coordinates of the 167,013 unique peaks as keys instead of gene names. When searching, the user does not need to specify the exact coordinates of the peaks; scfind will automatically find all peaks inside an interval to use as query. Compared to the full expression matrix, scfind achieves compression ratios of 10-300, and is more compact than other file formats (Fig 1b, S1). Both the relative compression and the absolute size of the index depends on the sparsity of the dataset, with the highest compression for MOCA where >95% of the expression matrix consists of zeros (Fig 1c). Since an index for 100,000 cells takes up no more than ∼150 MB of disk space, scfind makes it possible to analyze very large datasets on a standard laptop. Since the four largest scRNA-seq datasets contain only ∼10^3^ reads/cell, the range of expression values is limited, allowing them to be well approximated with only two bits/cell for the lossy compression (Fig 1d). The compression scheme used by scfind is not just memory efficient, but also fast; it typically takes less than a minute for each tissue to create an index (Fig S2).

To identify the cells that match a query, scfind decompresses the strings associated with each key to retrieve the cells with non-zero expression. If cell labels have been provided, scfind will automatically group the cells and a hypergeometric test is used to determine if the number of cells found in each cell type is larger than expected by chance. Search times are well below one second, even for the MOCA dataset with ∼2 million cells, and they scale linearly both with the number of genes in the query and the number of cells in the reference (Fig 1e,f).

As an example, we consider the genes Il2ra, Ptprc, Il7r and Ctla4, which correspond to T cell surface markers commonly used in FACS experiments. For the TM FACS dataset the results show that T cells in the bone marrow is the most highly enriched group, and further inspection reveals that 22 of the 30 cells that express all four genes are T cells from different tissues, suggesting that this is the most relevant hit (Fig 2a).

**Figure 2:**
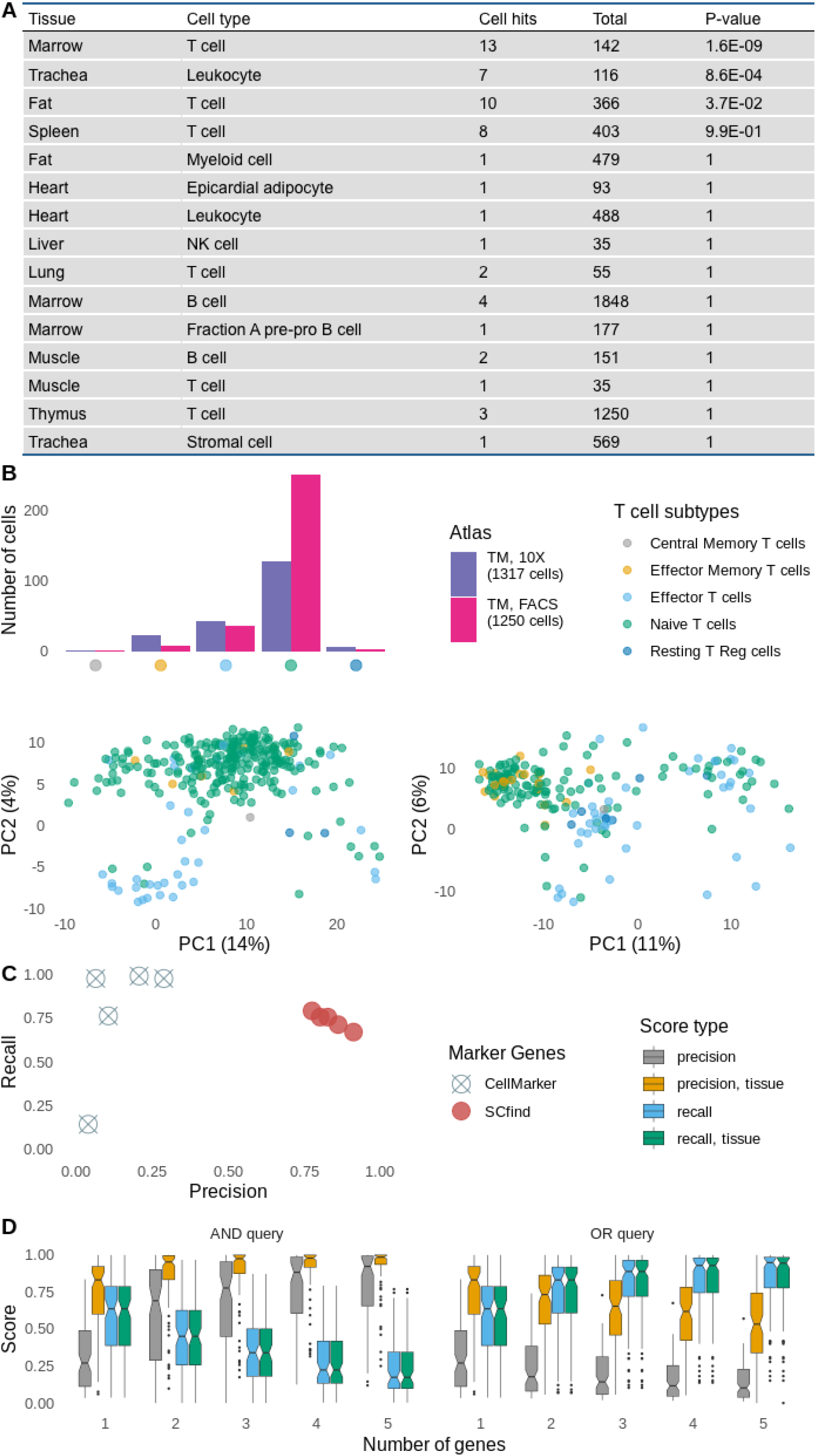
Basic search, *in silico* sorting and identification of marker genes. (a) Sample query showing the results for a search of the TM FACS dataset for Il2ra, Ptprc, Il7r and Ctla4. (b) Number of cells found for subsets of T-cells in the thymus based on combinations of Il2ra, Ptprc, Il7r and Ctla4 (top) and PCA projection of the T cells from the thymus shows good separation between naive T cells and resting T regulatory cells for both TM FACS (bottom, left) and 10X (bottom, right). (c) Evaluation of manually curated cardiomyocyte marker genes (Acta1, Actc1, Atp2a2, Myh6 and Nppa) compared to the five best based on F1 score for the TM FACS dataset. (d) Precision and recall values for combinations of 1-5 marker genes using AND or OR logic were calculated for all cell types in the TM FACS data. The boxplots show the spread of scores when compared either to all cell types or just the ones from the same tissue.

The default is to require that cells contain all keys in a query, but the user can specify for each key if it is to be used with OR or NOT logic. The logical operators make it possible to form complex queries and one application is to carry out *in silico* gating of cells, i.e. subsetting cell states of one cell type by their gene expression status by taking advantage of the fact that scfind returns individual cells matching a query. As an illustration, we considered the T cells from the thymus in the two TM datasets. Even though there are numerous subsets described in the literature, both datasets only contain a single cell type representing all T cells. Based on the expression of Il2ra, Ptprc, Il7r and Ctla4, we defined five non-overlapping sub-clusters as suggested by (Golubovskaya and Wu, 2016). The sub-clusters were detected in similar proportions in both datasets and they are similar to what has been reported elsewhere (Fig 2b, S3) (Belizário et al., 2016).

### Fast and flexible identification of marker genes

The ability to identify all cells expressing a gene in milliseconds makes it possible to exhaustively scan the entire transcriptome. That is, scfind can be used for optimization problems by evaluating all genes based on how well they perform for a specific task. One such application is to search for marker genes, i.e. genes that are present in one cell type and absent in all others. To evaluate gene g_i_ as a marker for cell type c_j_, we defined four quantities: true positives are cells that belong to c_j_ and express g_i_, false positives are cells from other cell types expressing g_i_, false negatives are cells from c_j_ that do not express g_i_, and true negatives are cells from other cell types that do not express g_i_. From these quantities precision, recall and F1 score can be computed to rank the genes. One can also combine multiple marker genes through AND or OR logic. Using AND to improve precision will typically result in reduced recall (and vice versa for OR), so combining markers will alter the trade-off between precision and recall, but it rarely improves the F1 score. To illustrate the marker genes search, we evaluate 5 markers for cardiomyocytes taken from the manually curated CellMarker database (Zhang et al., 2019) for the TM FACS dataset. Interestingly, these marker genes have moderate recall values but low precision values (Fig 2c). By contrast, scfind can identify marker genes that result in either high precision, recall, or F1 score (Fig. 2c, Table S1, S2). Considering all cell types, ranking all genes took only an average of 0.28 s for TM FACS, 0.18 s for TM 10X, and 0.74 s for MCA (Fig. S4), suggesting that an entire atlas can be ranked in minutes.

To compare how well marker genes perform across mouse cell types, we identified the best combinations of up to five marker genes in all three adult whole body atlases. Interestingly, the quality of the marker genes varies drastically across cell-types suggesting that some cell types are easier to distinguish than others. Closer inspection reveals that the cell types that perform poorly are typically ones that are found in multiple tissues, e.g. fibroblasts or immune cells. This is not necessarily unexpected since the differences between similar cell types from different tissues is expected to be relatively small. By contrast, cell types that are found in only one organ, e.g. hepatocytes or pancreatic beta cells, are easy to distinguish as they contain genes that are not expressed elsewhere. Consequently, when we modified the search to only compare against cell types from the same tissue, the scores increased (Fig. 2d, Table S1, S2).

By definition, the set of marker genes comprises the smallest number of genes that allows a cell type to be reliably identified. However, other criteria can be defined for selecting genes relevant for a cell type. For example, one can use the largest possible set of genes that are expressed above a threshold, and we define the maximal set of marker genes as those genes with recall>.9. Since recall is defined as TP/(TP + FN) the maximal marker set is independent of the other cell types in the atlas, unlike a definition based on precision or F1. Moreover, despite being selected based on their recall, maximal marker genes can achieve a high precision since very few cells from other cell types will match the query. The cardinality of the maximal marker gene set gives an indication of heterogeneity as a homogeneous cluster will have a larger number of genes. For example, the mean number of maximal markers for the TM FACS cell types is 238, but T cells, which we know have different subsets (Fig 2b) have only on average, 114 maximal markers (Table S4).

We can also use scfind to identify genes with similar expression patterns to a gene set specified by the user. The remaining genes are ranked based on how similar their expression pattern is to the query as quantified by the Jaccard index. The similarity search function can be useful if we have identified a set of good markers, but no antibodies are available. To find a suitable alternative we can search for genes with similar expression profile, restricted to the list of genes for which antibodies are available. As an example, we consider the five cardiomyocyte markers that we identified previously. Comparison with a list of 1,296 surface markers (Bausch-Fluck et al., 2015) reveals that none of them can be found on the surface, but scfind’s similarity search function returns five other markers for which antibodies are available (Fig S5).

### Identification of housekeeping genes and cell type specific genes

In addition to identifying sets of genes that are specific to a cell type, we can also evaluate how many different cell types express a specific gene. We define a gene as present if it is found in at least *max(*10, 0.1*N_i_)* cells, where *N_i_* is the total number of cells of type *j*. Consistent with previous studies based on bulk RNA-seq (Ju et al., 2013) we find a bimodal pattern such that many genes are either widely expressed or found in only a few cell types (Fig 3a). The number of cell types where a gene is expressed is strongly correlated between the different atlases (Spearman’s rho = .63 on average), suggesting that the findings are robust despite the differences in sequencing depth and cell type annotations.

**Figure 3:**
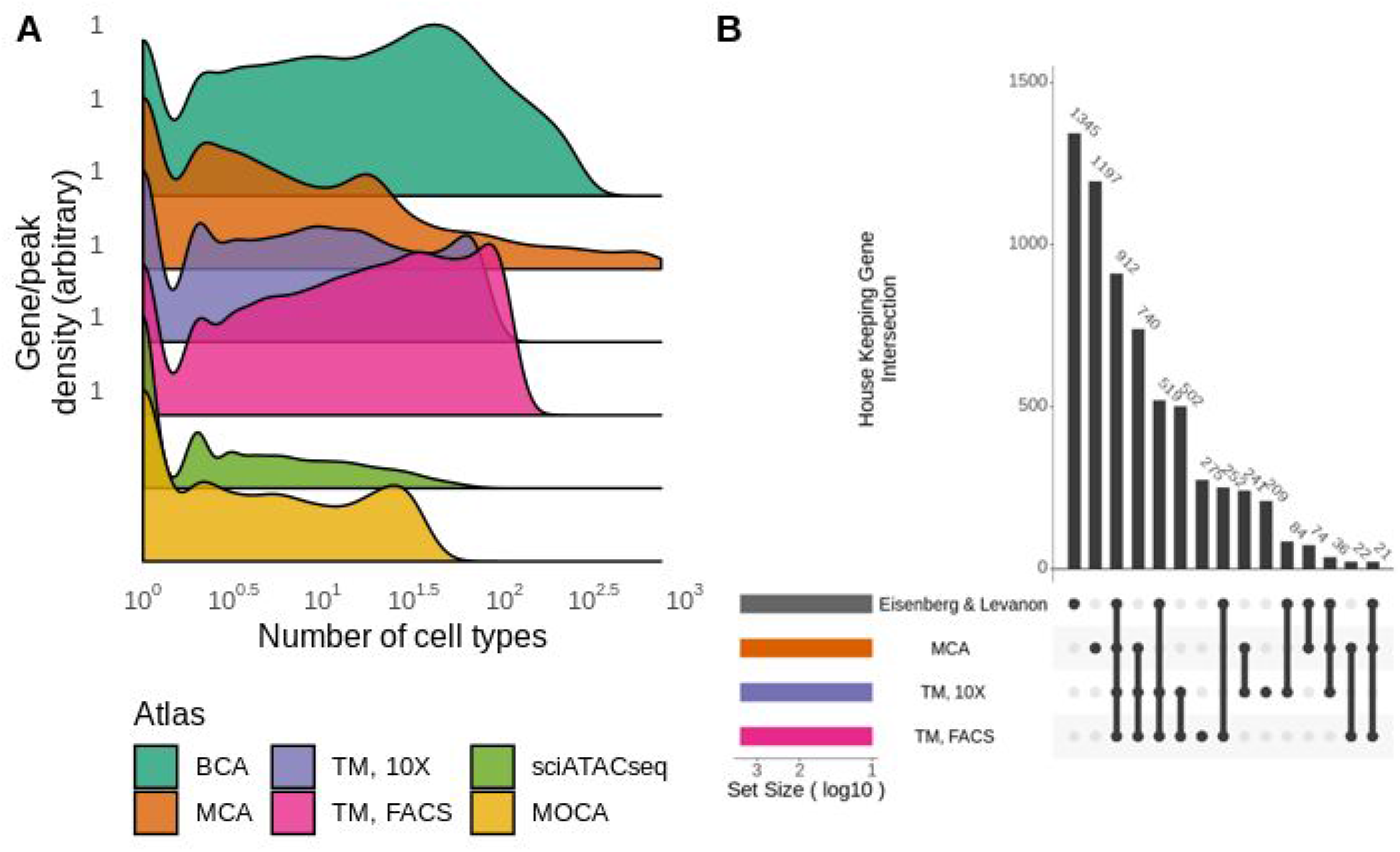
Evaluating cell type specificity. (a) Histogram showing the distribution of cell type specificities for genes and peaks from the six atlases. (b) Upset plot showing the overlap of 3,144 housekeeping genes from the literature and three mouse cell atlates.

One class of genes of particular interest are those expressed across most or all cell types, a.k.a. housekeeping genes. The most widely used definition of housekeeping genes is a list of 3,505 genes that was obtained from bulk RNA-seq data from several tissues (Eisenberg and Levanon, 2013). We compared the 3,144 genes from this list which were present in the MCA and the two TM datasets with the same number of widely expressed genes from each of the three atlases. This revealed a high degree of similarity, as 912 genes were found on all four lists (Fig 3b, Table S3). An additional 740 genes were found in all three atlases but not in the literature curated list, suggesting that there are a substantial number of ubiquitously expressed genes that are not traditionally considered as housekeeping genes. Moreover, 1,345 of the genes from the literature were not identified as widely expressed in any of the three atlases, implying that the results can differ substantially depending on whether bulk or single cell data is used. Since the ranking of the genes takes only ∼1 second/cell type (Fig S6), it will be straightforward to continuously refine lists of housekeeping genes as more data becomes available.

At the other end of the spectrum, it is also interesting to identify genes that are expressed in only one or a few cell types. We found 4,966 genes present in only one or two cell types in all three atlases (Table S3). Even though this result is influenced by the sequencing depth and the granularity used for the clustering, the implication is that thousands of genes are highly specialized and only found in a limited number of cell types. Closer inspection of the cell types containing the largest number of specific genes shows that they are mainly found in hepatocytes, brain and testis while many of the immune related cell types have few or no unique genes (Table S5).

### Frequent pattern growth algorithm ensures that long queries return meaningful results

One challenge in using queries involving even a moderate number (>5) of terms for sparsely sequenced datasets is that it is very likely that an empty set of cells will be returned. To ensure that meaningful results are returned without requiring the user to manually modify the query through trial and error, scfind features a query optimization routine. This procedure identifies subsets of the original query terms, hereafter referred to as subqueries that are guaranteed to return non-empty sets of cells. Since the number of possible subqueries may be very large, evaluating all possible combinations is intractable. Instead, scfind uses a strategy inspired by the frequent pattern growth algorithm (Han et al., 2004) to identify subqueries that return many cells. To help the user decide which subquery to use, they are ranked by a score inspired by the term frequency-inverse document frequency (TF-IDF) metric (Sparck Jones, 1972).

To illustrate the subquery optimization we consider the genes Acta1, Actc1, Atp2a2, Myh6, Nppa, Ryr2, Tnnc1 and Tpm, which can distinguish cardiac muscle cells in the heart (Piccini et al., 2015). A search of the TM FACS dataset returns no cells, but the subquery optimization ranks the query involving Actc1, Myh6 and Nppa as the best one. A search for these three genes shows heart cardiac muscle cells as the only significantly enriched cell type (Fig. 4a). As a more realistic and biologically meaningful positive control queries, we considered the GO database (The Gene Ontology Consortium, 2017) where each term is associated with a manually curated list of genes. Although some terms are unlikely to be cell type specific (e.g. “DNA binding”), for some terms it is reasonable to expect that they should be enriched for one or more cell types. For example, after using the subquery optimization we find that the best matches for “lung saccule development” (GO:0060430) has the highest enrichment in several different cell types found in the lung for the MCA, MOCA, TM 10X and TM FACS datasets. We queried all terms containing between 5 and 25 genes against the three mouse atlases, and report the p-values of all cell types for the best subquery (Fig 4b, S7, S8). Biclustering of the resulting matrix reveals that the rows, corresponding to the cell types, are not grouped by tissue. Instead, functionally similar cell types, e.g. immune cells, are grouped together, regardless of their tissue of origin. By contrast, the columns which represent the GO terms, form a hierarchy which corresponds well to the GO annotation evidenced by the fact that the parent categories are not mixed. This result is reassuring since the GO terms are hierarchical and can be represented as a tree structure with terms holding a similar meaning grouped together. Another important observation is that the subquery optimization takes <10 seconds even for queries involving up to 25 genes (Fig 4c). Taken together, we have demonstrated that subquery optimization can yield biologically meaningful results in a timely manner.

**Figure 4:**
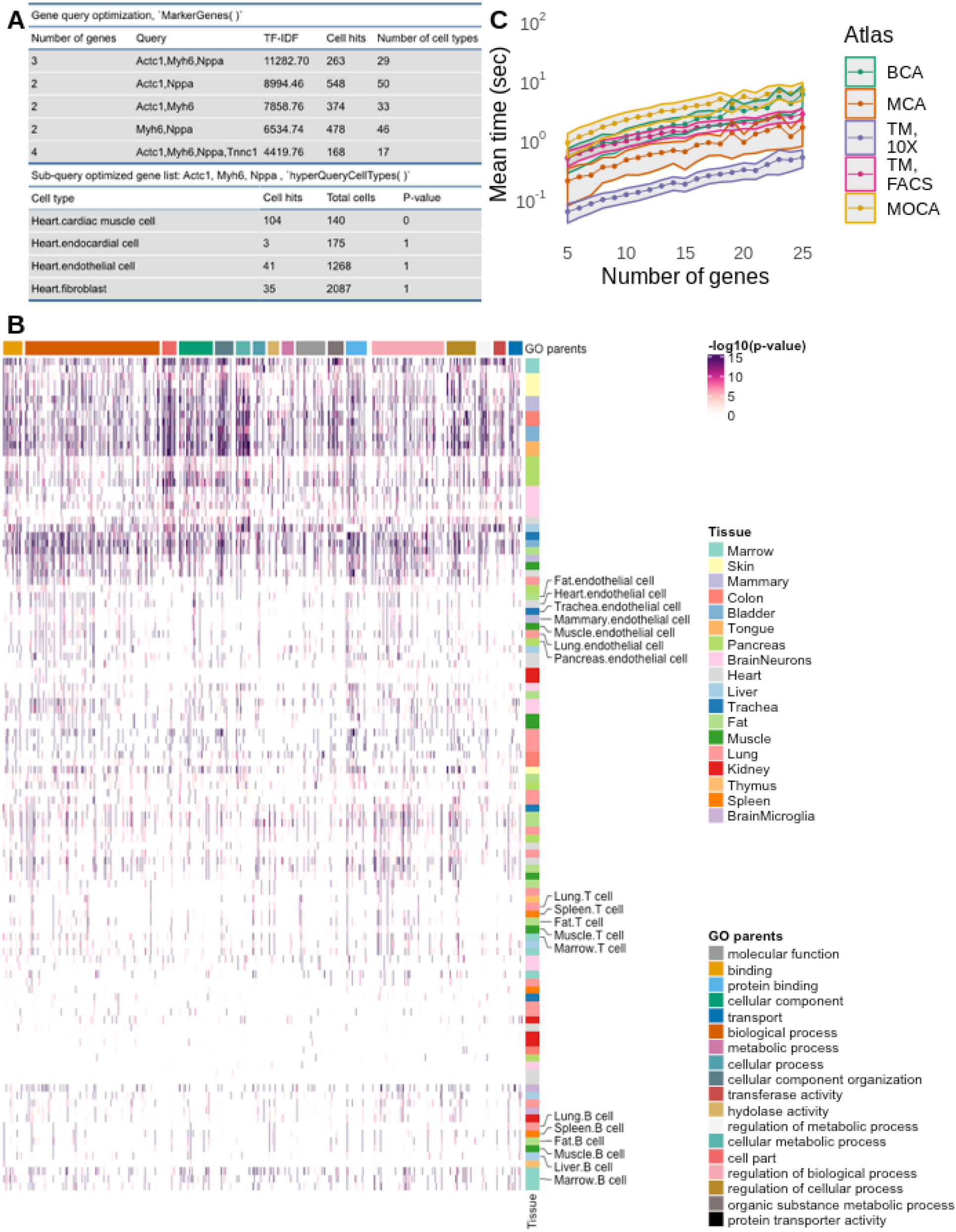
Subquery optimization. (a) Example for optimization of the query Acta1, Actc1, Atp2a2, Myh6, Nppa, Ryr2, Tnnc1 and Tpm for the TM FACS dataset (b) Heatmap showing the p-values for the best subquery for each GO term containing between 5 and 25 genes for the TM FACS data. Rows represent cell types and columns represent GO terms. (c) Mean run times for subquery optimization for lists from the GO annotation containing between 5 and 25 genes. The bands indicate standard errors.

### Queries involving biomedical keywords

One of the central purposes of a cell atlas is to facilitate the identification of cells or cell types associated with a specific query. We have demonstrated that scfind provides this functionality given a query formed by a list of genes. Although such queries are convenient for researchers with expertise in genetics and genomics, it restricts the number of people that can use the cell atlas. To allow for a much wider range of queries, scfind leverages the knowledge that has been accumulated over decades in the form of abstracts from the biomedical research literature stored in NIH’s PubMed database (Sayers et al., 2019). The PubTator (Wei et al., 2013) resource has parsed all PubMed abstracts up until 2018 and extracted gene names, genetic variants, disease names and chemicals for each PubMed ID. We assume that if a keyword, i.e. variant, disease or chemical, is mentioned in the same abstract as a gene, then the gene is relevant for queries involving that keyword. Based on this logic we have created dictionaries where each keyword is associated with a list of three or more genes sorted by how often they co-occur. In total, the dictionaries map >300,000 keywords to gene lists (Table 1a).

**Table 1:**
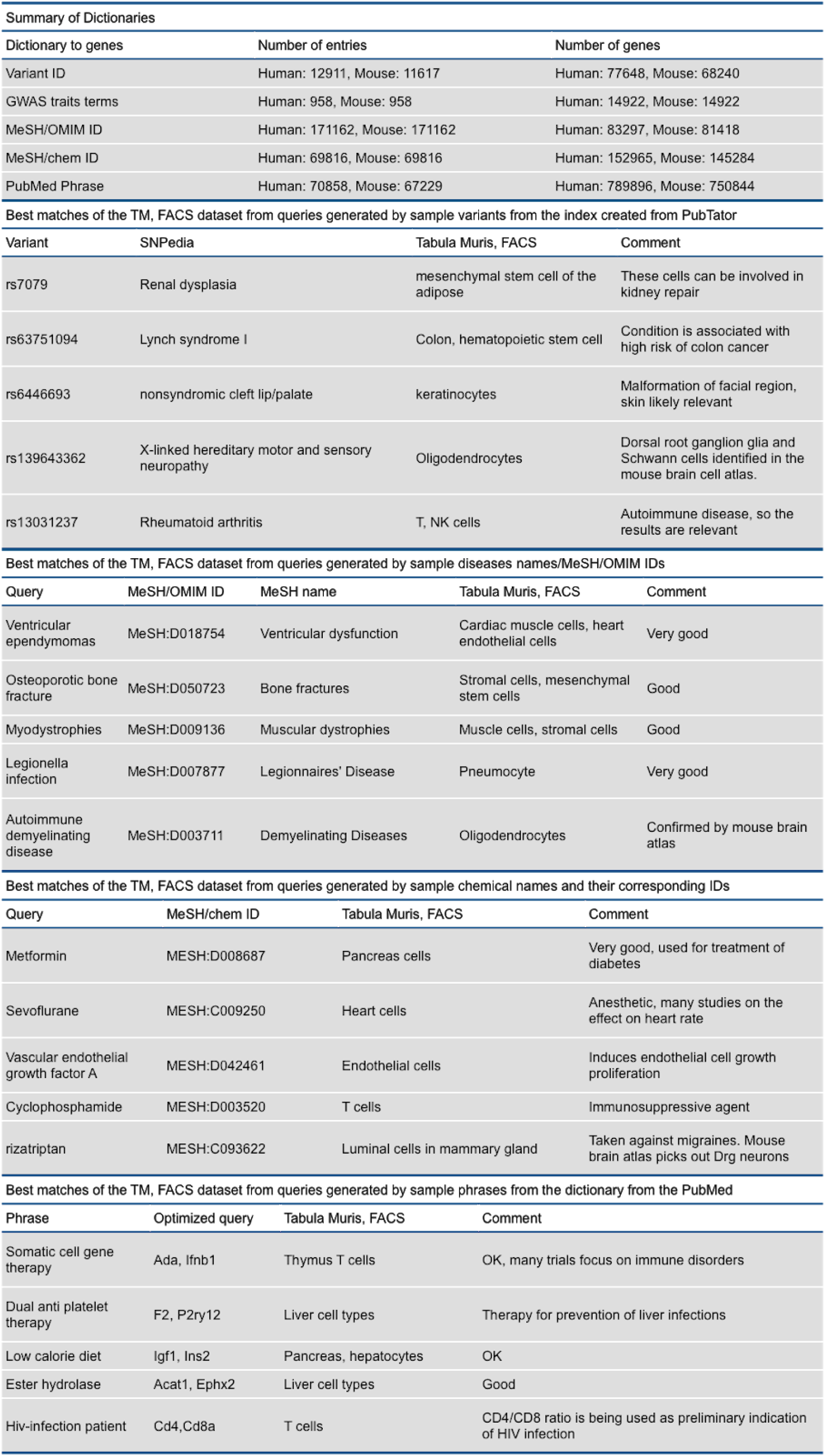
Queries involving keywords mapped to gene lists. (a) Summary statistics of the number of keywords ang genes for different categories. (b-d) Five variants (b), diseases (c) and chemicals (d) from the index created from PubTator. The association from SNPedia is listed as well as the best hits from the TM FACS dataset. (e) Five randomly selected phrases from the dictionary generated from the PubMed Phrases resource. The top hits from the TM FACS dataset are also listed.

The variant dictionary makes it possible to combine text mining and expression analysis to identify the cell type where a variant is most likely to have an impact. This approach is not new (Manica et al., 2019; Pers et al., 2015; Raychaudhuri et al., 2009), but scfind is the first method specifically designed for single-cell data. We report the results for five variants when queried against the TM FACS data (Table 1b). For the most part the results are informative, pointing to cell types that are likely to be relevant given what we know about the human phenotypes associated with each variant. An intriguing result is the pathogenic variant rs139643362 which is related to X-linked hereditary motor and sensory neuropathy (Cariaso and Lennon, 2012). The only match from the brain cell types are oligodendrocytes, a result confirmed when searching the mouse brain atlas which returns matches for the glia and Schwann cells from the dorsal root ganglion. This finding is supported in the literature where the disease has been implicated by lack of myelination (Scherer and Kleopa, 2012).

Many diseases go by a diverse range of names and PubTator addresses this issue by mapping each disease name to categories from Medical Subject Headings (MeSH) and Online Mendelian Inheritance in Man (OMIM). The map used by scfind relates 171,162 disease names to 6,639 disease categories. Each disease category is then associated with a gene list, and we find matches for 5,253 and 5,157 categories in human and mouse, respectively. Again, we provide five example queries to illustrate this feature (Table 1c). For example, a search for “Legionella infection” returns cell types from the lung as the top hits. Considering the etiology of Legionnaire’s disease, a bacterial infection causing pneumonia, this is clearly a relevant result.

Similarly, we have generated a map relating 69,816 chemicals to 69,812 categories defined by MeSH IDs and Chemical Entities of Biological Interest (Hastings et al., 2013). For human, 12,130 categories could be mapped to gene names, and for mouse 11,511 categories. This allows searches for drugs, hormones, toxins, etc and the results for five searches are reported here (Table 1d). To allow for word and phrase searches other than disease names and chemicals (that are common in the biomedical literature), scfind includes a dictionary, based on PubMed Phrases (Kim et al., 2018). The human dictionary contains 70,858 phrases and the mouse dictionary contains 67,229 phrases and can be used in the same way as the PubTator dictionaries (Table 1e).

The PubMed phrases expands the number of possible queries, but there are still significant limitations as the user is required to enter the precise keywords found in the dictionaries. To allow for greater flexibility, scfind employs a strategy from natural language processing, word2vec (Mikolov et al., 2013) to map queries that are not found in the dictionary to the existing keywords (Fig. 5a). The idea behind word2vec is to embed words in a vector space which makes it possible to match words and phrases based on similarity of their associated vectors. The neural network carrying out the embedding needs to be trained on a large corpus of text, and scfind uses one based on PubMed abstracts (Pyysalo et al., 2013).

**Figure 5:**
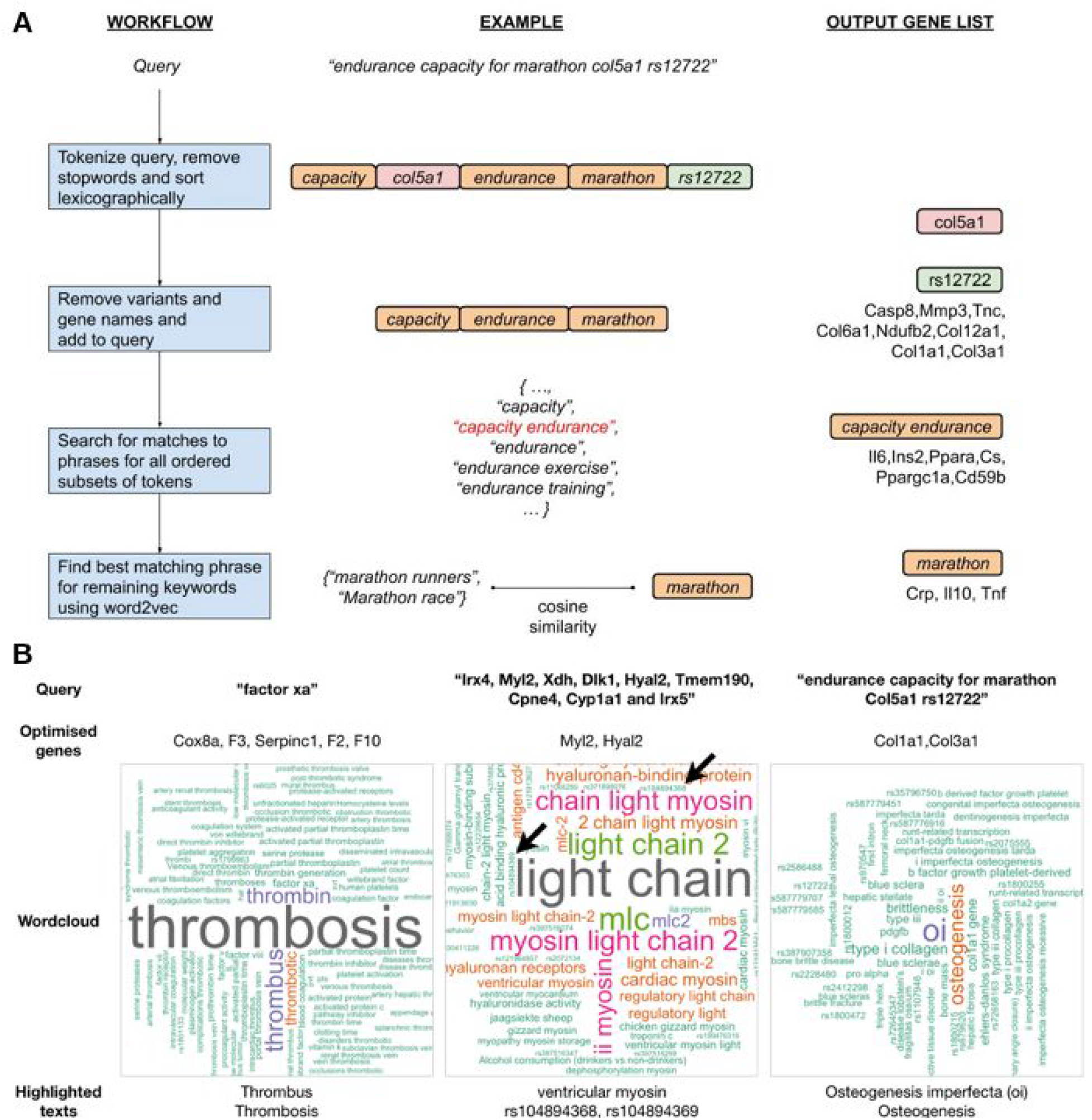
Free text searches and visualization of results. (a) Flow chart showing how a query is processed by by scfind. (b) Examples of word cloud associations following query optimization. Arrows in the middle panel show variant names (see main text).

Addition of the basic gene queries to the logic operators, the subquery optimization, the dictionaries based on PubMed abstracts, and the word2vec functionality, results in a versatile search engine that can identify cell types corresponding to complex queries at speeds that allow for an interactive workflow. When given a complex query, scfind first identifies gene names and keywords that exists in one of the dictionaries. Any remaining terms are then mapped using word2vec to the nearest keyword, and the gene names obtained are then used as a query (Fig 5a). The default is to use a strict query where the genes are combined using AND, but they can also be combined in a more permissive OR fashion.

The dictionaries mapping keywords to gene names can be inverted to provide a set of keywords associated with each gene. The inverted dictionaries are useful for interpreting a list of genes, and an intuitive way to visualize the results is through a word cloud where the size of each word is related to how frequently it is associated with the gene set. One application is to make sure that the gene list obtained following subquery optimization still represents the user’s original intent. As an example, we consider a query for Irx4, Myl2, Xdh, Dlk1, Hyal2, Tmem190, Cpne4, Cyp1a1 and Irx5 which are markers discriminating between atrial and ventricular heart tissue (Piccini et al., 2015). Subquery optimization for the TM FACS datasets suggests Hyal2 and Myl2 as the best query, and it shows cardiomyocytes and heart epithelial cells as the most enriched cell types. In the resulting word cloud (Fig. 5c), we note the prominence of the term “ventricular” which suggests that the sub query still reflects one of the key properties of the original gene list. Moreover, the word cloud contains several variants, eg rs104894368 and rs104894369 that have been associated with hypertrophic cardiomyopathy (Alfares et al., 2015; Flavigny et al., 1998), thus providing novel suggestions and interpretations to the user. It is worth emphasizing that the word cloud feature is not dependent on single cell data and can be used with any gene list.

### Application to single cell ATAC-seq data identifies cell type specific enhancers

Super enhancers, sometimes referred to as stretch enhancers (Hnisz et al., 2013; Parker et al., 2013), are defined as regions >3 kb that harbor a larger number of enhancer sites. Super enhancers are important for gene regulation and they have been extensively studied, but previous genome-wide studies have largely been based on bulk data. From dbSuper (Khan and Zhang, 2016) we obtained 4,328 super enhancer loci from eight tissues. For each super enhancer, we identified the most significantly enriched cell types by searching for overlapping peaks using an OR query, thereby assigning 2,726 loci to a unique cell type (Fig 6a, Table S6). This includes several known loci, e.g. Hnf4a, Nr5a2, Ppara and Rxra in hepatocytes (Joo et al., 2019) and Nr4a1 in monocytes (Thomas et al., 2016), that were correctly identified.

**Figure 6:**
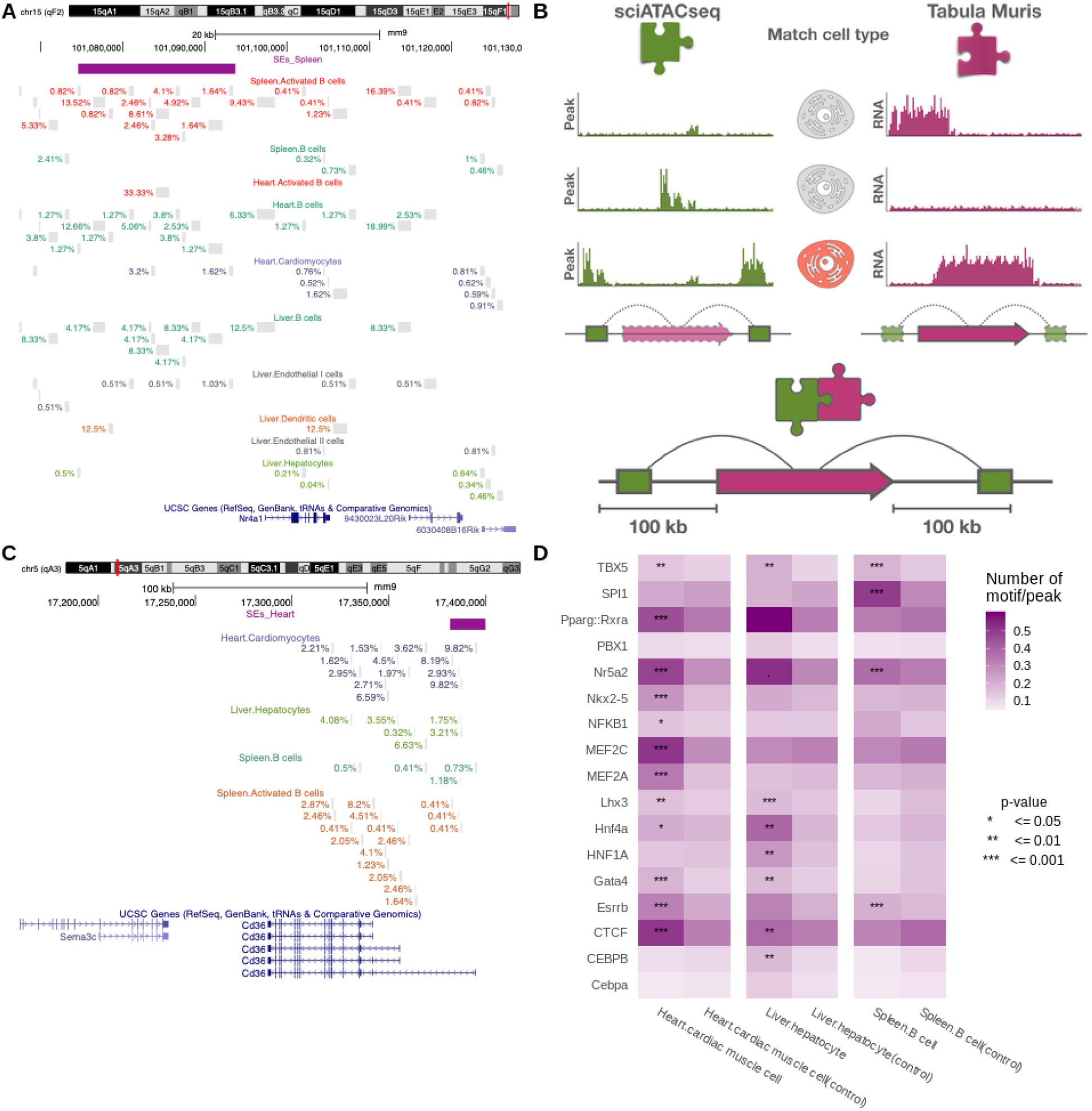
Determining cell type specificity of distal enhancers using scATAC-seq data. (a) UCSC genome browser screenshot showing the Nr4a1 locus which has a super enhancer in pro-B cells and monocytes. Grey marks represent open chromatin and the numbers show the fraction of cells containing each peak. (b) Schematic illustration of how scATAC-seq and scRNA-seq atalases are combined to identify cell type specific distal open chromatin loci and their targets (c) UCSC genome browser screenshot of the Cd36 locus showing open chromatin in several cell types (d) Motif enrichment in putative distal enhancers that are specific to cardiac muscle cells, hepatocyte and B cells.

One of the biggest challenges in studying gene regulation in metazoans is to identify the targets of distal enhancer elements. As enhancer targets cannot be predicted from sequence alone, information both about chromatin and gene expression is required (Kleftogiannis et al., 2016). Since proximity can be a useful guide, we hypothesize that open chromatin peaks that are only found in one cell type are likely to target nearby genes that are specifically expressed in the same cell type. Based on this rationale, we first searched the TM FACS atlas for genes that are highly expressed in only one cell type for each tissue. For each of those genes, we searched the sciATAC-seq atlas for peaks that were unique to the same cell type and within 100 kb of one the start sites (Fig 6c). This procedure identified 15,583 open chromatin-gene pairs from seven tissues (Table S7). We found support in the literature for several of the identified candidates, e.g. an enhancer upstream of Cd36 which is specific to cardiomyocytes (Fig 6b) (Khan and Zhang, 2016) and one upstream of Myh6 in cardiac muscle cells (Dickel et al., 2016). As a validation, we searched for enriched transcription factor binding motifs in the putative enhancers for three cell types where more than X loci were identified. This analysis revealed an enrichment of motifs bound by Mef2 in cardiomyocytes, while binding sites for Hnf1a and Hnf4a were enriched in hepatocytes (Fig. 6d).

Scfind can also be applied to multi-omics datasets where more than one assay is used for each individual cell. As an illustration, we created an index for the sciCAR data which provides both transcriptome and open chromatin information for 13,395 cells from the mouse kidney (Cao et al., 2018). The combined index has a total of 28,669 genes and 249,382 peaks, and queries can include both types of keys. We modified the marker gene search method to carry out an AND query for each gene along with each of the peaks found within 100 kb of its TSS. This allowed us to identify putative enhancer-gene pairs that are specific to each cell type. As an example, we identified 112 genes and 442 putative enhancers for the paranephric body adipocytes (Fig 7). Although the data is sparse and noisy, our analysis of adipocytes identifies an open chromatin region ∼60 kbps upstream of the Car12, a gene which has been shown to be expressed in adipose tissues and associated with type 2 diabetes (Zhong et al., 2010).

**Figure 7:**
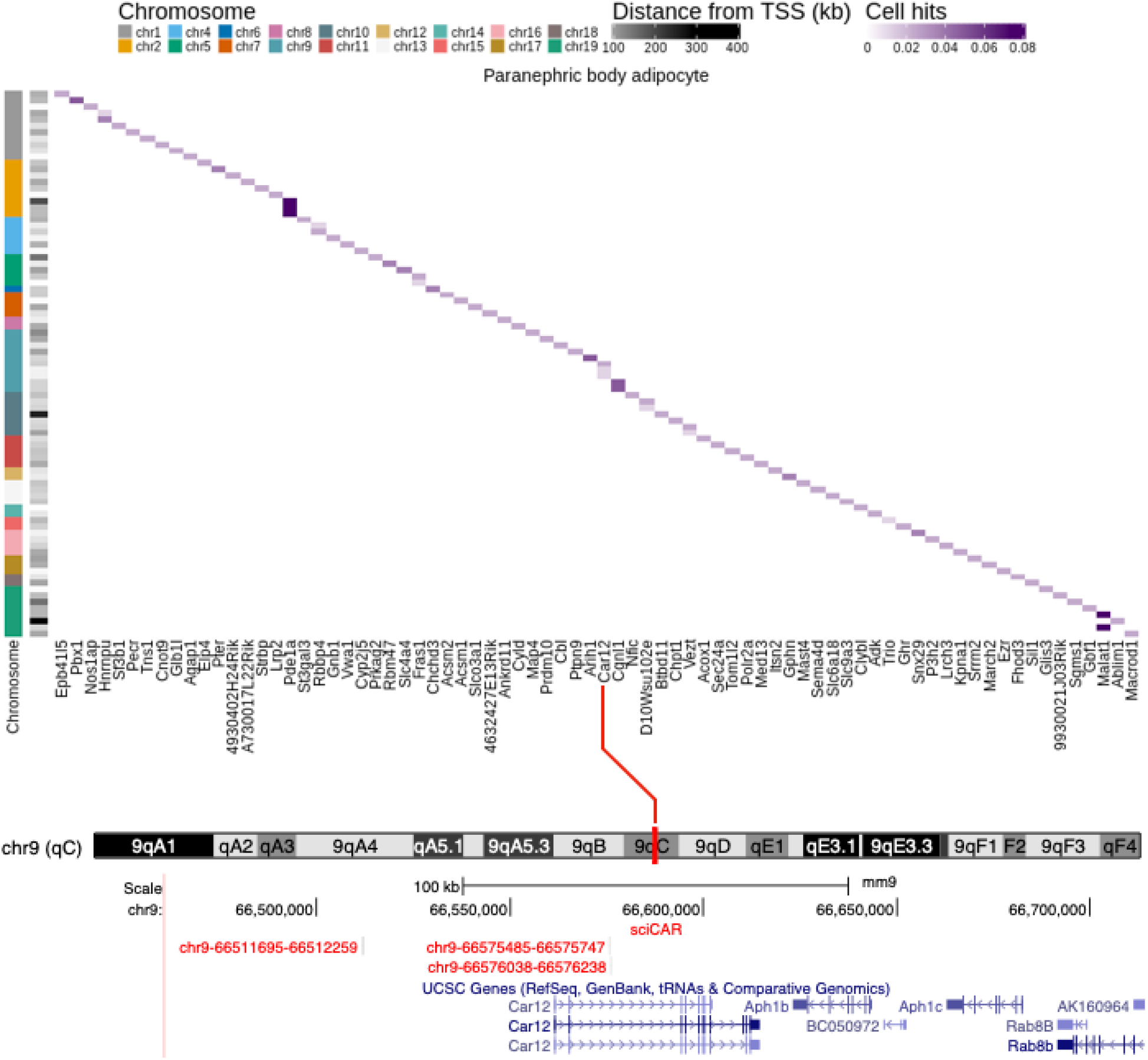
scfind for multi-omics data. (*top*) Heatmap showing the fraction of paranephric body adipocytes that have both open chromatin (row) and expressed gene (column). (*bottom*) UCSC screenshot of the Car12 locus with the putative enhancer marked in red.

## Discussion

As single-cell datasets grow exponentially (Svensson et al., 2018), there is a need for computational tools to allow for fast and efficient analysis. We have presented scfind, a method for searching large collections of single cell data to identify sets of cells that correspond to complex criteria. The main advantage of scfind is that it can carry out complex queries for datasets containing millions of cells in less than a second, while requiring only modest hardware. It has been shown that sub-second response times are required for interactive use of Internet search engines (Brutlag et al., 2008), suggesting that scfind can support real-time exploration of cell atlases. A web interface with a few pre-calculated indexes is available at https://scfind.sanger.ac.uk. Unlike other databases of scRNA-seq data, scfind retains information about individual cells and not just pre-defined clusters. Since scfind is freely available as an R package as part of Bioconductor with source code available under the MIT licence at https://github.com/hemberg-lab/scfind, it is easy for users to build their own references and carry out searches based on groupings other than the ones provided by the original authors. Because of its performance, scfind can be used for tasks that otherwise would have been prohibitive, e.g. running searches of all genes to be able to identify marker genes, similar genes, housekeeping genes, and cell type specific genes. These global evaluations can be carried out in seconds, making it possible to continuously update information on expression profiles, even for very large collections of data.

Our subquery optimization routine combined with the PubMed based dictionaries makes it possible to resolve arbitrary queries, ensuring that scfind can be used by people without expertise in genetics and genomics. The searches are difficult to evaluate quantitatively, but our tests suggest that they frequently provide reasonable results. Nevertheless, there are several ways by which results could be improved. The way in which the information is extracted is not very sophisticated, and associations are based solely on co-occurrence without considering the context. For example, an article may incorrectly put forward evidence that a specific gene is *not* important for a disease. Although we cannot rule out that some searches will be confounded, we conjecture that the impact of this may be limited since positive result bias is well documented in biomedical science i (Callaham et al., 1998; Fanelli, 2010). More advanced methods for processing natural language (Cambria and White, 2014) will make it possible to further exploit this knowledge when interpreting and analyzing high-throughput datasets.

To search a cell atlas, three main components are required; (i) experimental data, (ii) clustering and biological annotation and (iii) a search algorithm, and for results to be meaningful, both the data and the computational methods need to be of high quality. The work presented here falls under the third category. However, most of the testing has been carried out using mouse datasets, even though much of the information that is used to relate queries to gene lists is based on human studies, and this may explain some of the discrepancies. Furthermore, scfind suffers from the well-known issue of garbage in-garbage out, and even if the datasets are of high quality and accurately annotated, the algorithm may return nonsense. For example, for some of the keywords in our dictionary, e.g. “Los Alamos”, “DNA content”, and “prospective longitudinal study”, it is difficult to establish what cell types constitute a good match. Yet, they map to gene lists and a search is likely to identify significantly enriched cell types. Another scenario where scfind may fail to produce meaningful results is if the relevant cell type is not present in the index. For example, Usher syndrome is a genetic disorder that affects vision and hearing. A search against the TM FACS dataset returns pancreatic B and D cells as the most significant hits, most likely due to the fact that there are no cell types from the eyes or ears in the atlas.

The primary use case for scfind is to query large collections of previously annotated single-cell datasets to identify the group of cells that provide the best match. To extrapolate the most data out of scfind, it would be advantageous to connect it to large collections of datasets, e.g. Panglao (Franzén et al., 2019), scRNASeqDB (Cao et al., 2017), SCPortalen (Abugessaisa et al., 2018), or the EBI Single Cell Expression Atlas (Athar et al., 2019). Scfind has been designed to be easy to integrate at the back-end of these portals. As maintaining such databases requires substantial effort and resources, it is our ambition that scfind will be adapted by others. As a second use case, scfind can facilitate the annotation of newly collected datasets. Once the cells have been grouped through unsupervised clustering, determining the biological significance of each of the clusters is typically a difficult and time-consuming process (Kiselev et al., 2019). It typically requires expert knowledge and the researcher must search the literature to match marker genes with known pathways, processes or cell types. Scfind has the potential to speed up this process in several ways. For example, the methods for marker gene identification will make it possible to quickly identify relevant gene sets and the search functions allow the user to find the cluster that best corresponds to a gene set that has been derived from other experiments from the literature.

## Author contributions

MH conceived of the project and supervised the research. JL, NP, VK and MH contributed to the code, JL, NP and MH analyzed the data, JL and MH wrote the manuscript with input from NP and VK.

## Acknowledgements

We would like to thank members of the Hemberg group, Yong Liu, Tobias Bergmann and Anaise Meziani for assisting with beta-testing of the software and Luz Garcia-Alonso, Vanessa Jane Hall, Alessandro Ori, Sarah Teichmann and Roser Vento for feedback on the manuscript. We would like to thank Jana Eliasova for assistance with figure 1a.

## Methods

### Compression and decompression

Compression is carried out separately for each row (typically representing a gene or a peak), *i*, and cell-type, *c*, and it is split into two steps. First, we use the Elias-Fano coding (Vigna, 2013) to store the array containing the indexes of cells with non-zero elements. The Elias-Fano code uses two bit strings, *H_ic_* and *L_ic_* to store an array of ascending integers. The sparsity of the array automatically determines how each values is stored in *H_ic_* and *L_ic_* to provide asymptotically optimal compression. The total number of cells, *n_c_,* and the number of cells with non-zero expression, *n_ic_*, are also stored to allow for decoding. Second, for all cells the non-zero expression values, *g_ij_*, are fit to a log-normal distribution and the mean, *m_ic_,* and the variance, *v_ic_,* are stored. Given a user-specified number of bits, *b*, each value *g_ij_* is assigned to one of the 2 quantiles. The bit vectors are then concatenated and stored as *B_ic_*. The bit strings *B_ic_*, *H_ic_* and *L_ic_* along with *n_ic_, m_ic_,* and *v_ic_* are stored as entries in a hash table with the row label as key. Since each cell-type is stored separately, it is easy to filter results by excluding cell-types, tissues or experiments.

When decoding, the indexes of the non-zero cells are first obtained from *H_ic_* and *L_ic_*. Since the decoded lists are sorted, intersection can be carried out in linear time. By decoding the lists with the smallest number of cells first, the search is further sped up. An approximation of the original expression value, *h_ij_*, can be obtained by identifying the midpoint of each quantile for the log-normal distribution with mean *m_ic_* and variance *v_ic_*.

### Compression ratios and file sizes

The compression ratios in Figure 1b were calculated using the R command `obj_size`, comparing the size of the original expression matrix and the compressed scfind index. The index sizes reported in Fig 1c correspond to the file-sizes when saved to disk.

### Search times

For Fig 1e,f we used the `searchMarkers` function to search for cells expressing the randomly selected genes. For the comparison, we used the `counts` function applied to each of the `SingleCellExperiment` objects used to represent the tissues from the TM FACS dataset in order to obtain the count matrix. To create SingleCellExperiment, Seurat and Loom objects, `SingleCellExperiment` function of the SingleCellExperiment package, `CreateSeuratObject` function of the Seurat package and `build_loom` function of the SCopeLoomR package are used respectively. The search times in Fig 1e were calculated as the average of 100 queries involving random sets of genes. For both panels we used rejection sampling to ensure that only queries returning a non-empty set of cells were considered. All analyses are performed on the Jupyterhub platform built with 100GB RAM and 4 Intel Xeon 2.50 GHz CPUs.

### Quantization accuracy

The Spearman correlation was computed using the built-in R function by comparing the quantized gene expression values, *q*, with the original values, *g*.

### Logical search operators

By adding ‘*’ in front of a gene name it will be used as OR, and by adding ‘-’ in front, it will be used as NOT. It is also possible to combine the two operators for a NOT OR query.

### Identification of T cell subsets

We defined the different non-overlapping subsets of T cells using combinations of Il2ra, Ptprc, Il7r and Ctla4 (Golubovskaya and Wu, 2016). The queries used were “-Il2ra, Ptprc, Il7r” (naive T cells), “Il2ra, -Il7r” (Effector T cells), “-*Il2ra, -*Ptprc, Il7r” (Effector memory T cells), “Il2ra, -Ptprc, Il7r” (Central memory T cells), and “Il2ra, Ptprc, Il7r, -Ctla4” (Resting T reg cells).

### Cell-type enrichment

To determine if cell-type *i* is enriched if *m_i_* cells are observed out of a total of *m* cells returned for the query, a hypergeometric test is used. Here, the total number of balls is given by *N*, the number of white balls in the urn is given by *n_i_*, the number of white balls drawn is given by *m_i_*, and the total number of draws is given by *m*. The reported p-value is further adjusted using the Benjamini-Hochberg procedure (Benjamini and Hochberg, 1995) to correct for multiple testing.

### Visualization

To aid in the visualization of the results, scfind calculates a UMAP (McInnes et al., 2018) projection of the cells during index construction. Cells that match a query are automatically identified in the UMAP projection. This feature can help the user identify patterns, e.g. sub-clusters, formed by the cells matching the query.

### Identification of cardiomyocyte marker genes

We ran the scfind commands `cellTypeMarkers` for the `cardiac muscle cell` cluster for the TM FACS datasets. The marker genes from the CellMarkers database were evaluated using the `evaluateCellTypeMarkers` command.

### Evaluation of precision and recall for marker gene search

When searching for marker genes, all genes are evaluated and ranked based on either precision, recall or F1 (default). When evaluating multiple markers in AND/OR fashion scfind employs a greedy strategy by considering the cells expressing all/at least one of the top *m* markers to calculate the precision and recall as described above.

### Identification of maximal marker gene sets

By default scfind uses 0.9, but it may need to be adjusted depending on sequencing depth.

### Similar genes

For the query terms, we first identify all corresponding cell indices, *C*. For each term *i* not in the query, we extract the list of indices, *D_i_*. The terms are then ranked based on the Jaccard similarity, |*C ∩ D_i_|/|C ∪ D_i_|*. For the cardiomyocyte example, we used the command `findSimilarGenes`.

### Frequent pattern growth search and TF-IDF scores

To identify subsets of the original query that are guaranteed to return cell hits we use a modified version of the frequent pattern growth (FPGrowth) algorithm. The FPGrowth algorithm is popular in data mining where it is used to identify groups of items that co-occur in transactions. In the context of scfind one may think of cells as transactions and genes or peaks as items.

We first modify the original query so that all terms are related by OR logic to retrieve all relevant cells. For each cell *c_j_*, an inverted cell index is constructed, containing a list of the subset of query terms found. The next step is to determine a minimum support threshold, *s*, i.e. the minimum number of cells required to consider a query. To speed up this process, we use a heuristic to ensure that the number of subqueries considered remains bounded. For each term in the query, *g_i_*, scfind estimates the overlap with other terms in the query, and for *n* terms, *s* is defined as the median of the *n(n-1)/2* values.

The FPGrowth algorithm takes a set of cells and builds a data structure called an FP-tree. In the FP-tree each node represents a term from the query, making it a more compact representation than a matrix as each node contains a counter to keep track of the number of cells where the term was present. To construct the tree, we use the inverted index to iterate across all cells. As the terms are inserted in order of decreasing frequency, the most widely used terms will be found near the root, and widely supported subqueries can be extracted by traversing the tree. To generate subqueries the tree is traversed depth first, but the search is interrupted when the value of a node is below *s*. Note that the heuristic is chosen to ensure that the number of subqueries is limited, even for very long queries (>50 genes). If a low number subqueries are returned initially, the threshold will be lowered to ensure that a sufficient number of subqueries is considered.

To help the user decide which subquery to choose, scfind ranks them using a score inspired by TF-IDF. The score is calculated for each element of the expression matrix as *t_ij_* = log(*e_ij_ /* Σ_i_*e_ij_*) - log(*N_i_ / N)*, where *e_ij_* is the quantized expression values, *N_i_* is the total number of cells with non-zero values for term *i* and *N* is the total number of cells in the reference. For a query involving *n* terms the score for each cell is given by 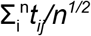.

To verify that the subquery optimization routine provides meaningful results we carried out a positive control experiment. For each cell type *c_j_* in each of the five scRNA-seq atlases we randomly selected five genes from the top 20 marker genes as defined by the F1 score. We calculated the top TF-IDF score and whether or not *c_j_* came up as the top ranked cell type for the best subquery. As a comparison, we also selected five genes from the top 100 marker genes. The results show that the former query results in significantly higher TF-IDF scores and is more likely to return the expected cell type as the top hit (Fig S9, S10, S11).

### Robustness analysis of TF-IDF search

We also investigated negative controls to test the robustness of scfind to changes in the expression matrix. We carried out an experiment whereby each element in the TM FACS data is multiplied by a random number which is uniformly distributed in the interval [.5, 2]. We then ran 1,000 queries containing gene lists from the GO database, and in 95% of cases the same cell type was ranked at the top for the best subquery as for the unperturbed data. Scfind is more sensitive to changes in the sparsity pattern and when a random selection of 10% of the elements in the expression matrix were either set to zero or given a non-zero value, the same cell type was chosen in 76% of cases. We also randomly permuted the cell labels for each tissue, and in this case only 51% of the queries returned the same cell type as the original query.

An important challenge in combining datasets into a cell atlas is to account for systematic technical differences between experiments, i.e. batch effects. Batch driven variation can be corrected *post-hoc* through computational means, but how these corrections can best be carried out remains an open problem. Several methods have been published in recent years (Tian et al., 2019), but some of these are incompatible with scfind as they do not operate on the expression matrix, but rather on another representation of the data, e.g. a selection of principal components or the distance matrix. The methods that do modify the expression matrix, e.g. Combat (Johnson et al., 2007) and Seurat v3 (Stuart et al., 2019), typically do not preserve zeros and may even introduce negative values. Consequently, the batch corrected expression matrix may violate one of the central assumptions that scfind makes about the data. An important constraint when applying batch correction methods prior to building an index is to ensure that zeros are preserved. To test the impact of batch correction methods directly, we implemented modified versions of Combat, Limma and Seurat v3 where only non-zero elements are adjusted and we used them to combine the two Tabula Muris datasets. We identified cell type markers for the merged dataset and for all three batch correction procedures we found that the top 500 genes were the same as for the TM FACS data in all cases. Taken together, we conclude that scaling and batch correction have limited impact on the results for scfind.

### Processing of PubTator and PubMed Phrases

A custom Julia script was used to build indexes mapping variants and phrases to mouse and human gene names. The indexes were also inverted to allow gene names to be mapped to phrases for the word cloud visualization. Processed dictionaries are available as additional files for the Homo sapiens and Mus musculus genes.

### Word2vec analysis

We used the Julia package `Word2Vec` for the calculation of cosine similarity between the tokens of the arbitrary query and the tokens of the similar phrases subsetted from the manually generated phrases to gene names dictionaries. To identify the best match phrase, the cosine distances between each pair of query-to-phrase token is calculated using the PubMed word2vec models. The distance of each phrase is averaged by the number of tokens. The mean distances are used to rank the best matched phrase.

### Wordcloud visualization

We use the R package `wordcloud`.

### Cell type specificity of super enhancers

For each locus from dbSUPER we used its mm9 coordinates to search for enriched cell types using an OR combination of the peaks at each loci. A locus is reported if it has a p-value<10^-10^, is found in at least 10 cells and >10% of the cells from that cell type.

### Cell type specific genes and peaks

A gene is considered cell type specific if it is found in at least 25 cells and enriched at a p-value <10^-5^ with respect to the other cell types in the tissue. For each cell type specific gene we considered all peaks within 100 kb of the 5’ most TSS. A peak is considered cell type specific if is present in at least 10 cells and at least 10% of the cells from one cell type. Moreover, we require that the peak does not meet the above criteria for any other cell type in the same tissue.

We manually selected motifs from the JASPAR 2018 Core collection (Khan et al., 2018). We then used the R packages `TFBStools` (Tan and Lenhard, 2016) and the `motifmatchr` package to identify motif instances in the cell type specific peaks.

### Analysis of sciCAR data

Indexes for the RNA-seq and ATAC-seq datasets were constructed separately and then merged according to corresponding cells. For each matrix which columns represent the cell of each cell type and rows represent both gene names and peaks loci, is organised as a `SingleCellExperiment` object. An scfind object is built by merging indexes of all cell types using the function `buildCellTypeIndex` and `mergeDataset`.

## Datasets used

### Mouse atlases

The data for the MCA was downloaded from https://figshare.com/s/865e694ad06d5857db4b, the Tabula Muris data from https://figshare.com/projects/Tabula_Muris_Transcriptomic_characterization_of_20_organs_and_tissues_from_Mus_musculus_at_single_cell_resolution/27733, the BCA data from http://linnarssonlab.org/data/, the MOCA data from https://oncoscape.v3.sttrcancer.org/atlas.gs.washington.edu.mouse.rna/downloads and the sciATAC-seq data from http://atlas.gs.washington.edu/mouse-atac/data/. For all datasets, custom R scripts were used to combine the expression and annotation files to generate `SingleCellExpression` objects (REF). The `SingleCellExpression` objects were used to build the scfind indexes.

### GO annotation

The GO annotation was downloaded from Ensembl Biomart website.

### PubTator, Pubmed Phrases and word2vec

The PubTator resources covering PubMed abstracts until 2018 was downloaded from ftp://ftp.ncbi.nlm.nih.gov/pub/lu/PubTator/ and the PubMed phrases dataset including abstracts until 2017 was downloaded from ftp://ftp.ncbi.nlm.nih.gov/pub/lu/PubMedPhrase/PubMed_Phrases.tar.gz. The word2vec model trained on PubMed abstracts was downloaded from http://bio.nlplab.org/.

### Super enhancers

We downloaded the .bed files corresponding to mm9 for Bone Marrow, Cerebellum, Heart, Kidney, Liver, Lung, Spleen and Thymus from the dbSUPER website. We also downloaded the .bed file containing hg19 coordinates for GM12878 cells.

## Code availability

The code for scfind is available at github.com/hemberg-lab/scfind and the code for generating the figures in this manuscript is available at https://github.com/hemberg-lab/scfind-paper-figures

## Data availability

The indices for the five atlas datasets can be downloaded from https://scfind.cog.sanger.ac.uk/browser.html?shared=indexes/.

## Supplementary tables

These are large and provided as separate files.

**Table S1:** Precision, recall and F1 scores for all cell types in the atlases considered.

**Table S2:** Information about the total number of marker genes and the precision and F1 scores that they provide for each cell type.

**Table S3:** Cell type specificity for the genes found in the MCA and the two Tabula Muris datasets.

**Table S4:** Number of maximal marker genes for each cell type in the MCA and the two Tabula Muris datasets.

**Table S5:** Number of cell type specific genes for each cell type in the MCA and the two Tabula Muris datasets.

**Table S6:** Cell type specificity of super enhancers

**Table S7:** Cell type specific enhancer-gene pairs

## Supplementary figures

**Figure S1:**
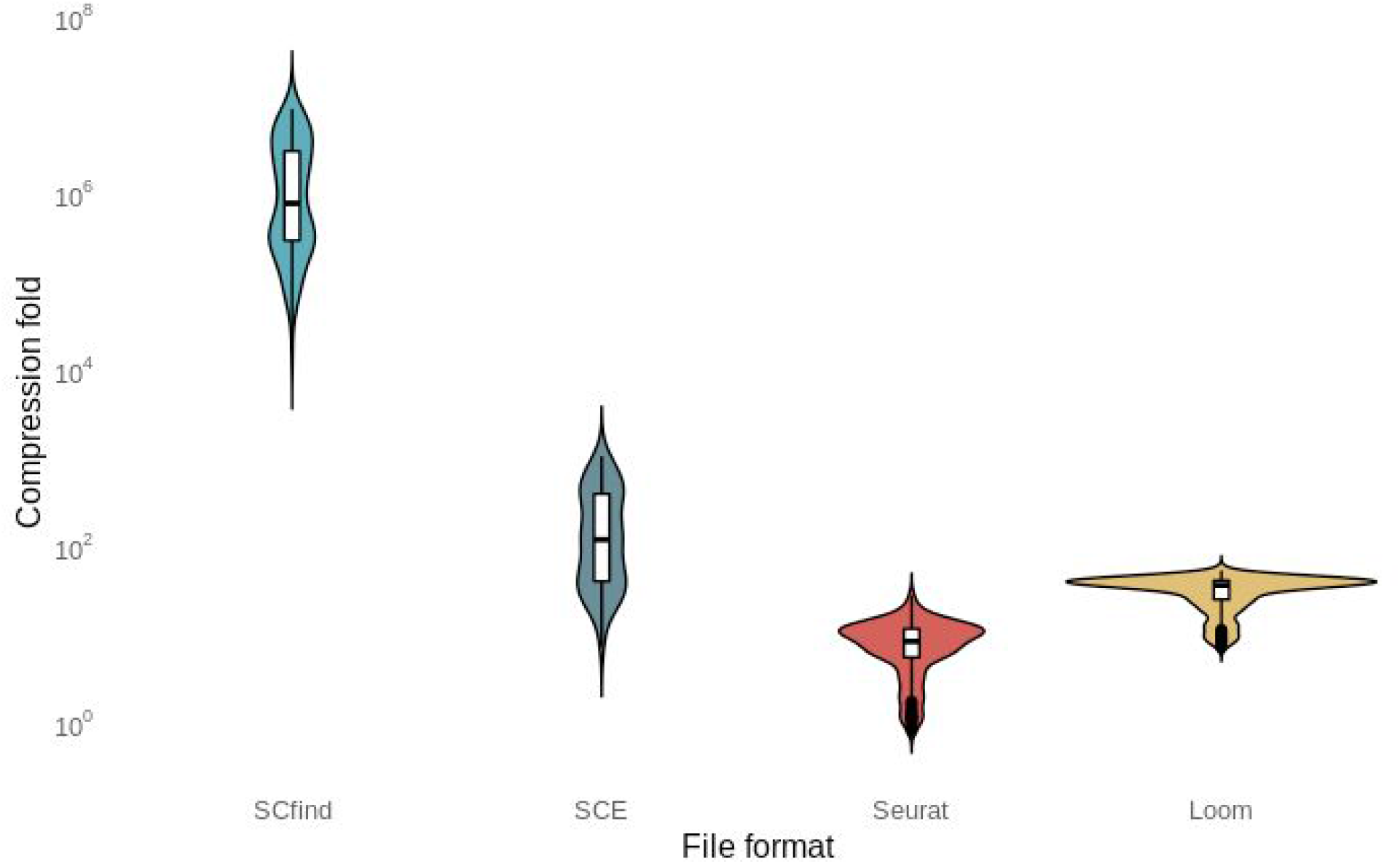
Compression ratios for different file formats, the violin plots represent the density of folds relative to the uncompressed expression matrix for all tissues in the six atlases in figure 1.

**Figure S2:**
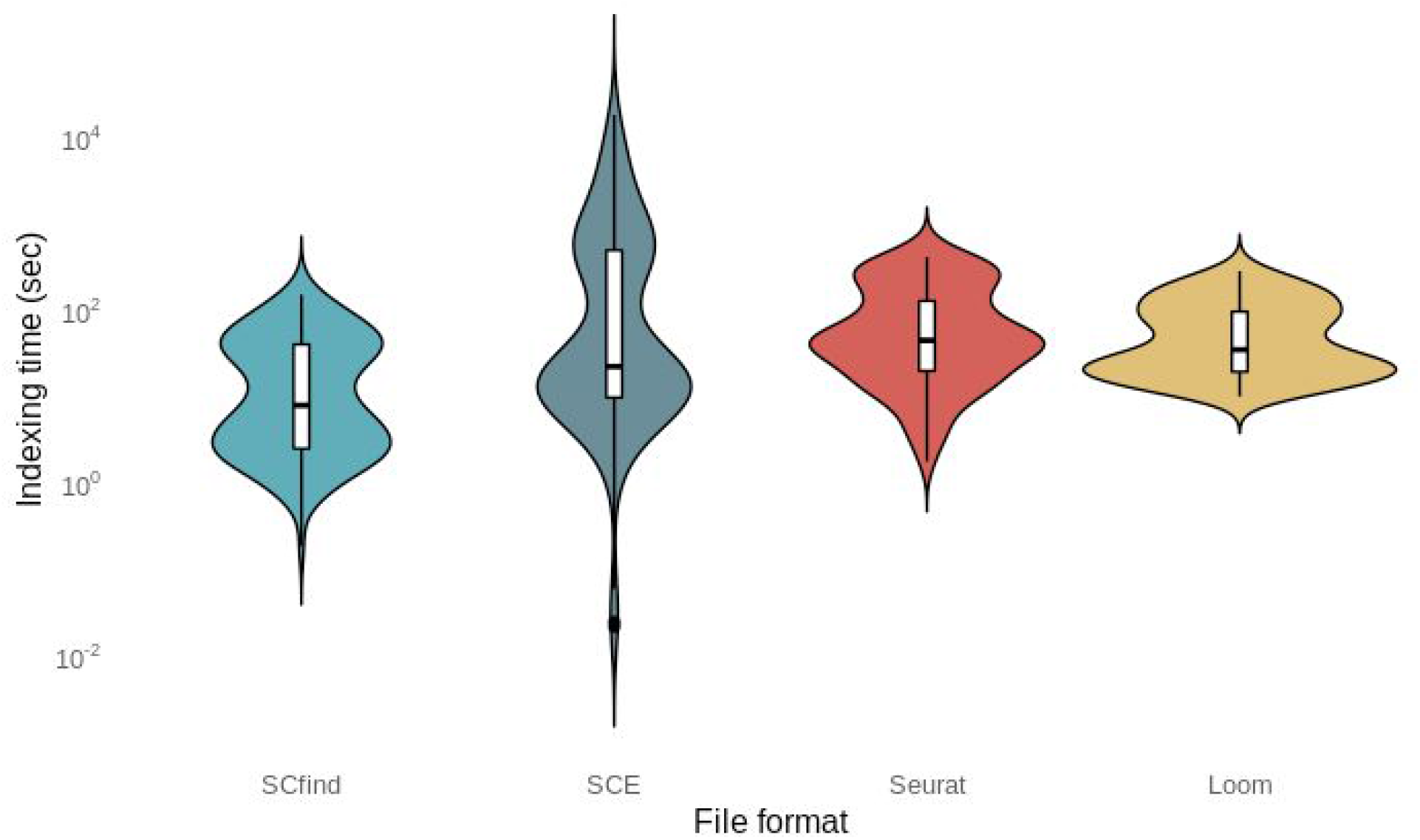
Build times for scfind indexes and other file formats, the violin plots represent the density of folds relative to the uncompressed expression matrix for all tissues in the six atlases in figure 1.

**Figure S3:**
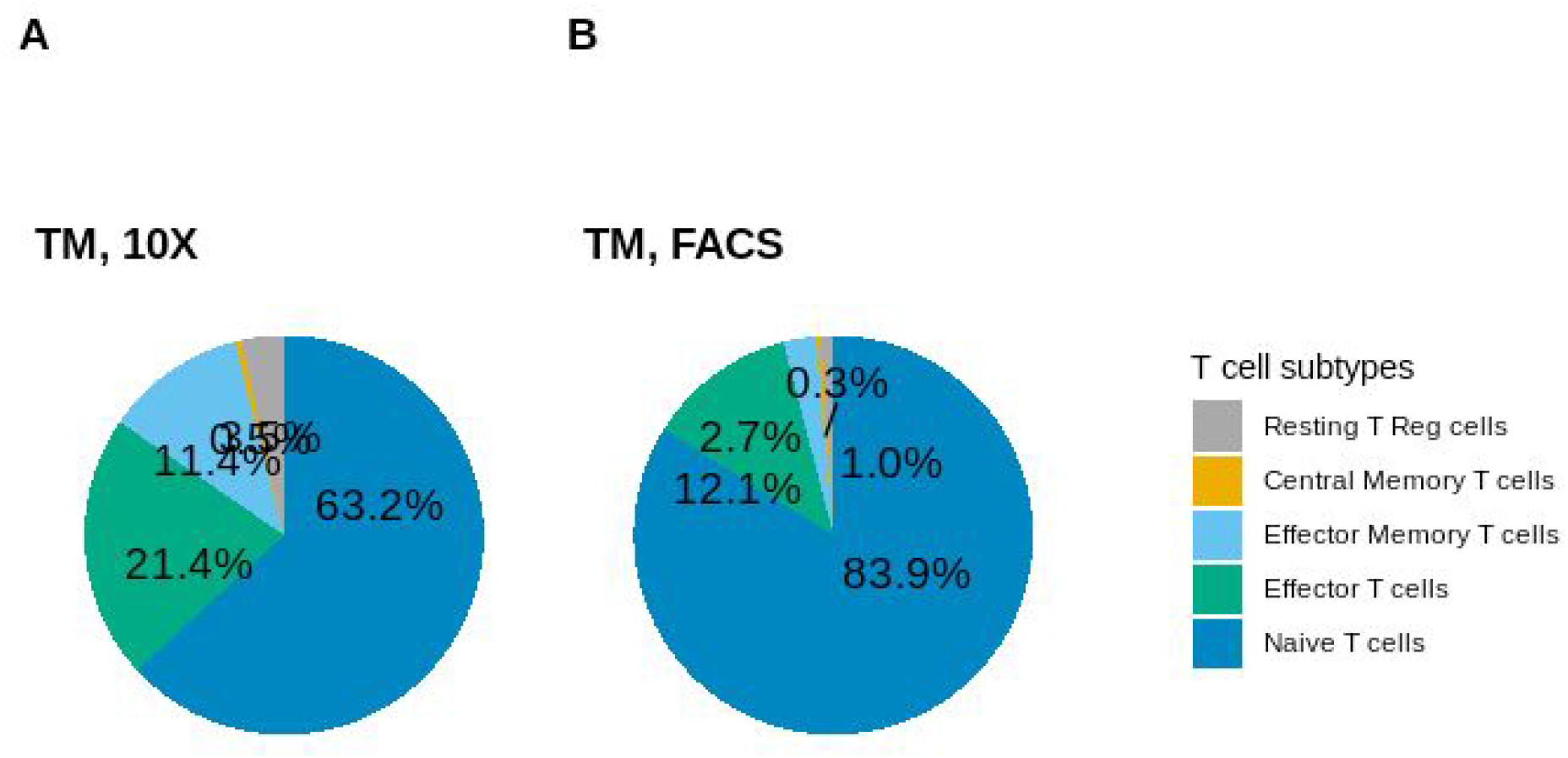
Percentage of T cell subtypes in thymus tissue of the Tabula Muris datasets

**Figure S4:**
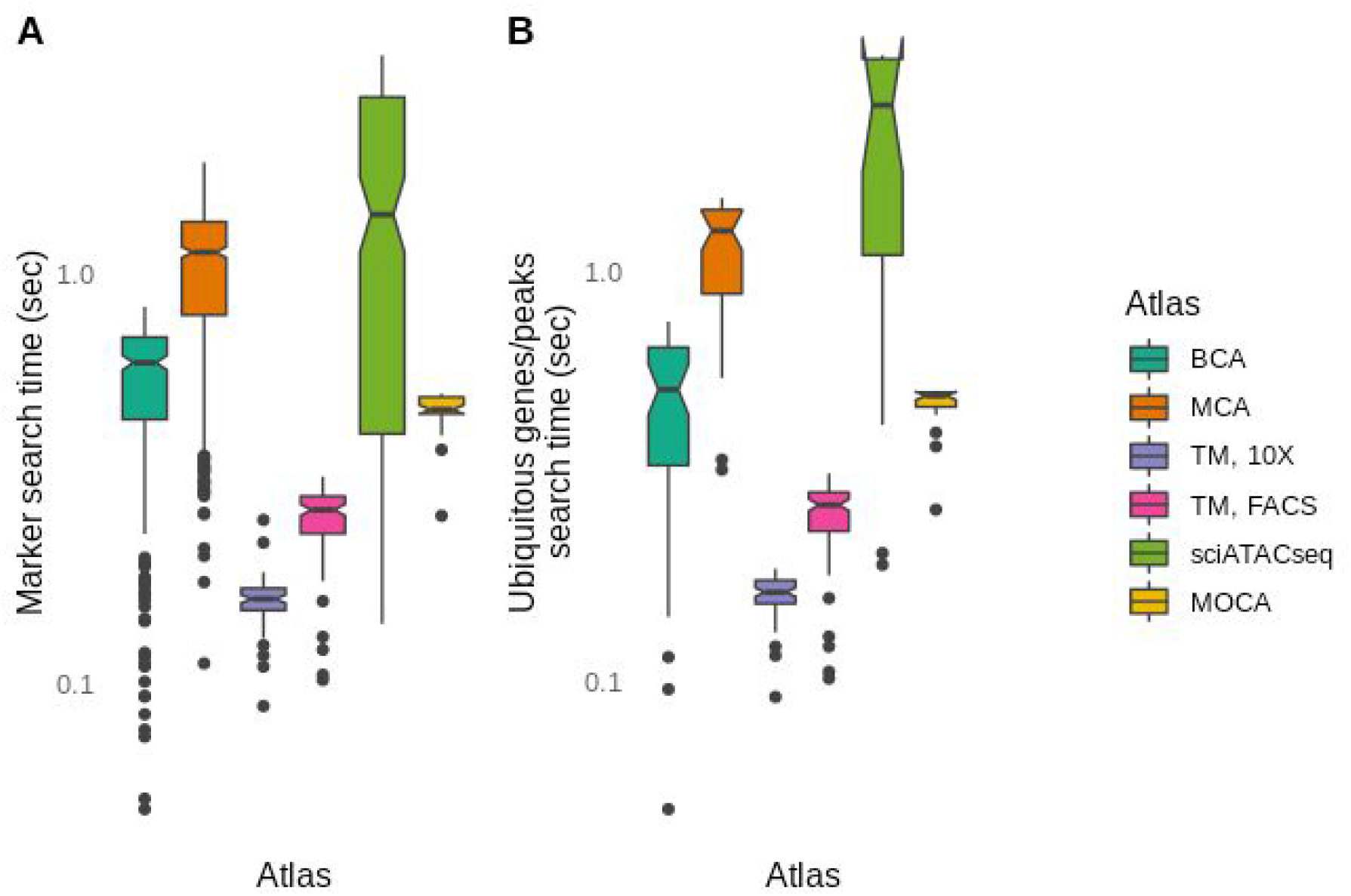
(a) Search times for marker genes/peaks and (b) evaluation of number of cell types where a gene is found.

**Figure S5:**
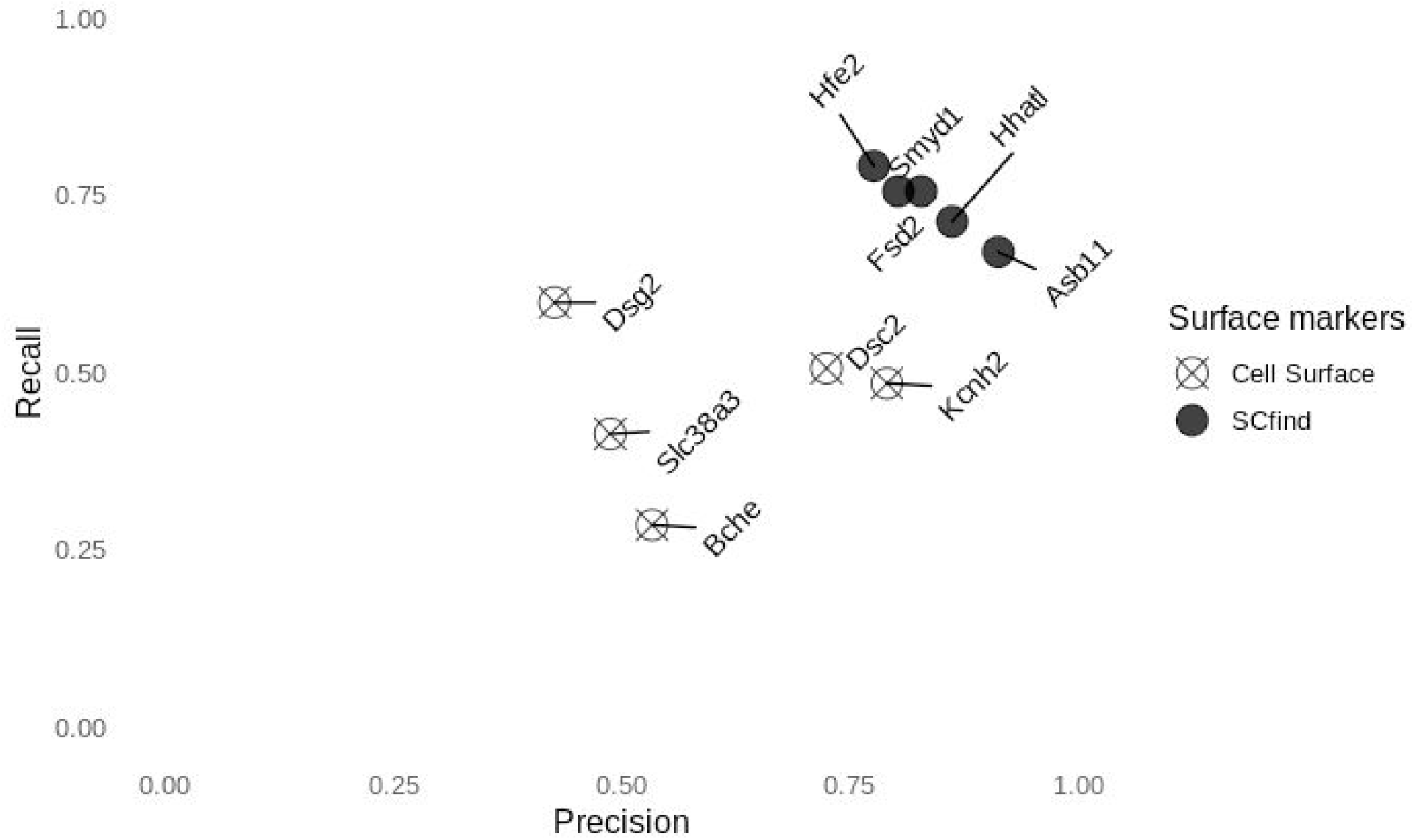
Precision and recall for the five best cardiomyocyte surface markers

**Figure S6:**
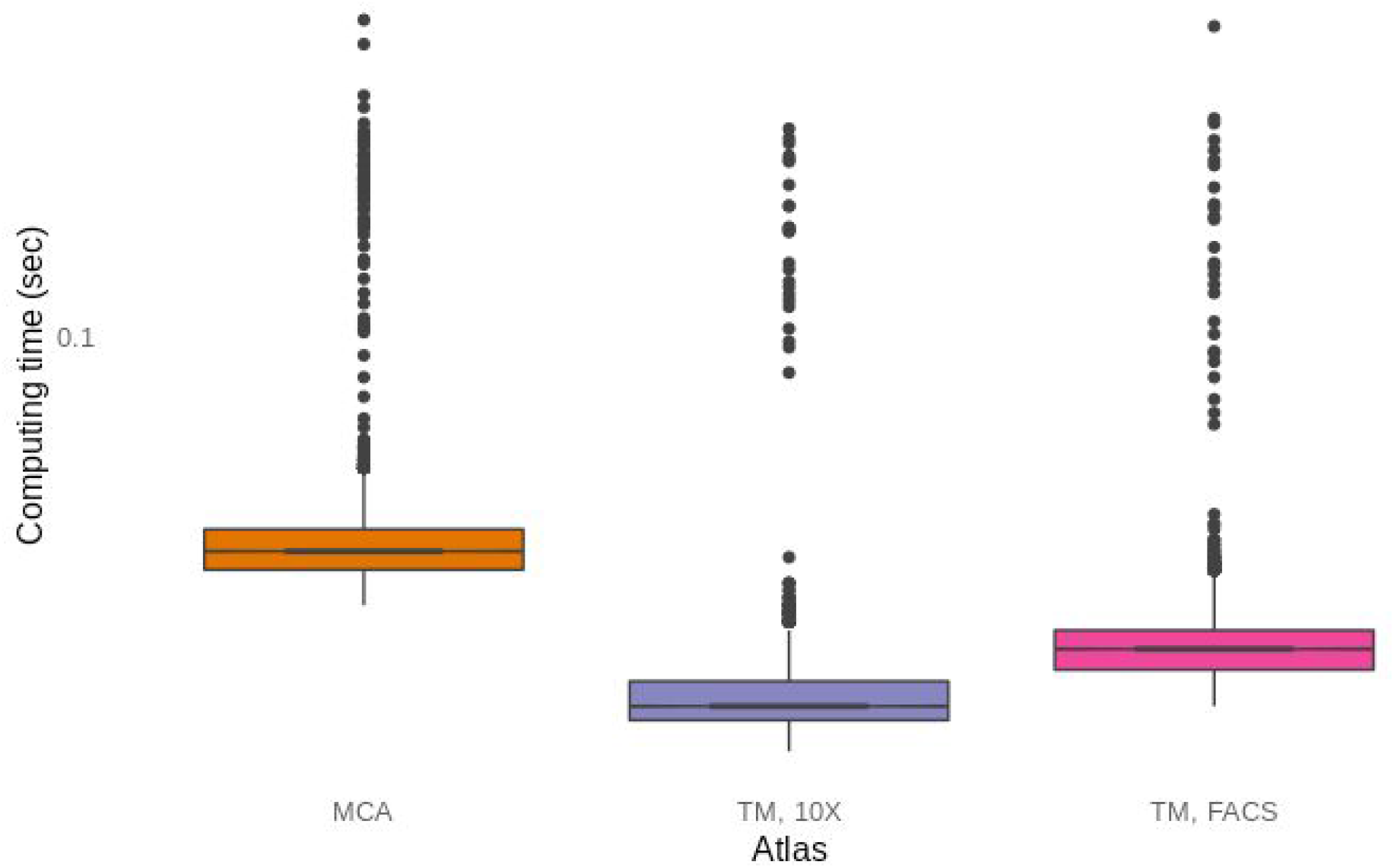
Times to calculate cell type specificity for the genes found in the MCA and the two Tabula Muris datasets.

**Figure S7:**
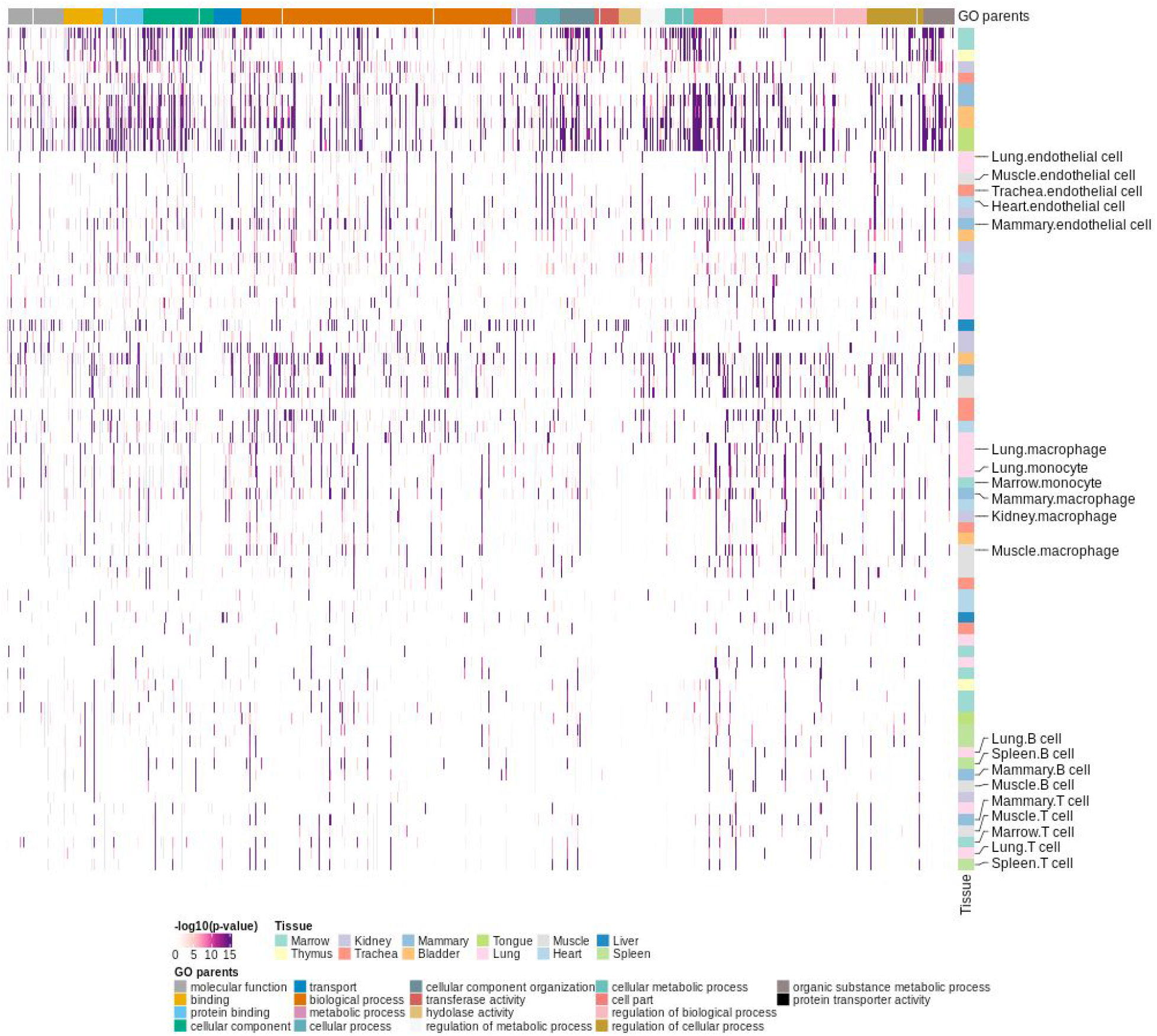
Heatmap showing the enrichment of cell types from the TM 10X data for GO terms with between 5 and 25 genes.

**Figure S8:**
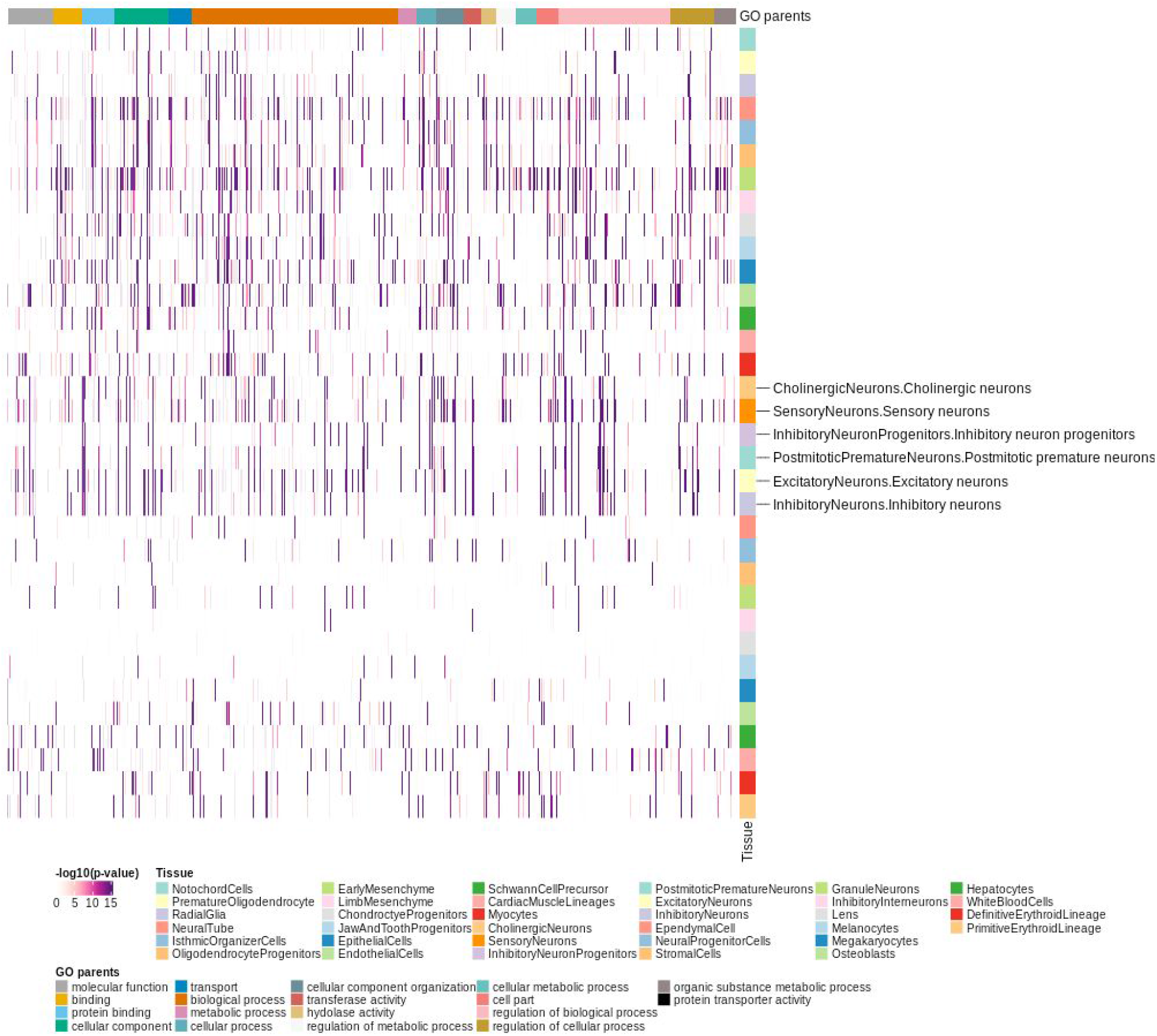
Heatmap showing the enrichment of cell types from the MOCA data for GO terms with between 5 and 25 genes.

**Figure S9:**
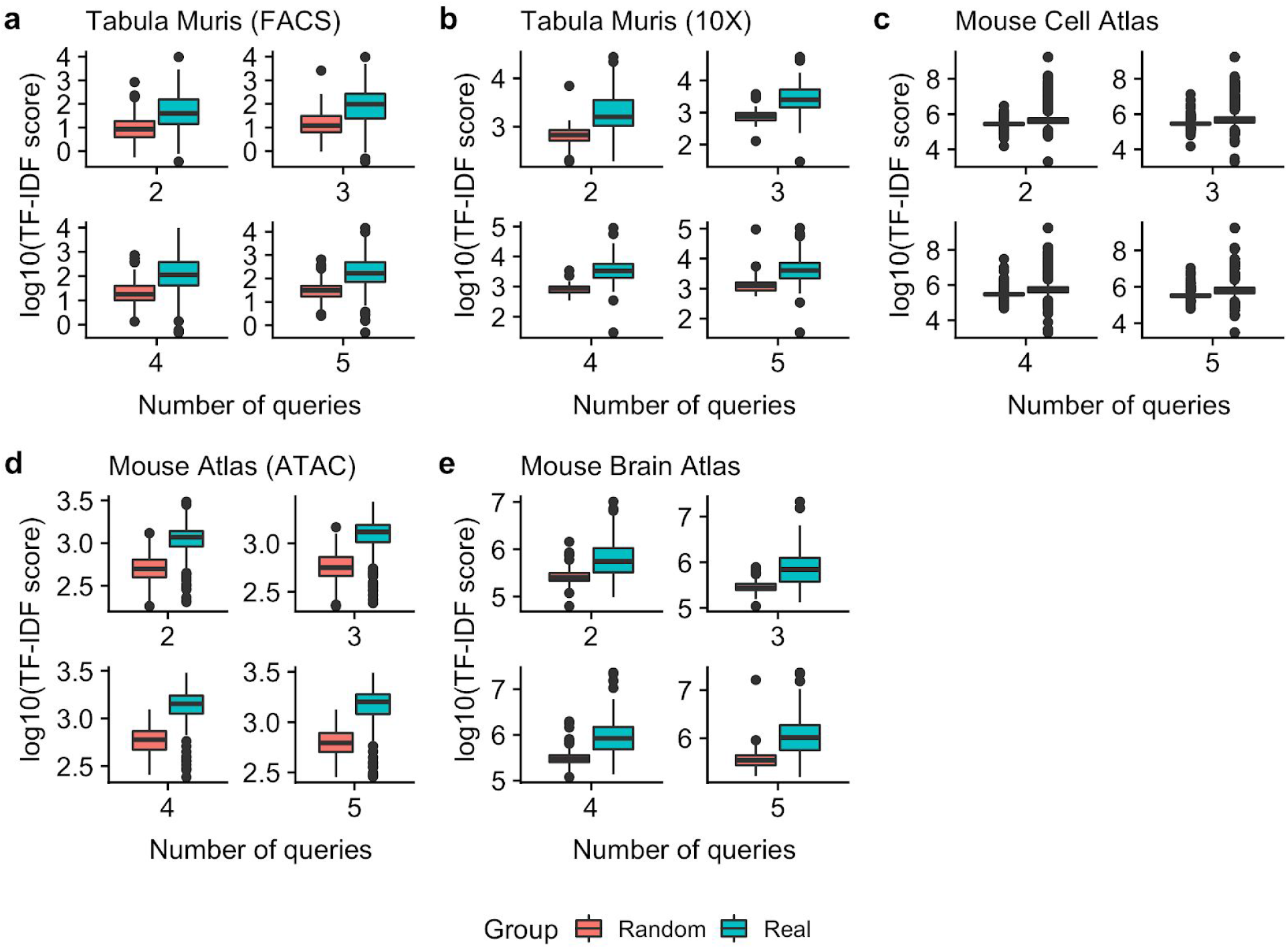
Comparison of TF-IDF score between queries based on the top 20 (Real) and top 100 (Random) marker genes for the (a) Tabula Muris (FACS), (b) Tabula Muris (10X), (c) MCA, and (d) Mouse Atlas (ATAC) datasets, (e) Mouse Brain Atlas. Fifty sets of real gene queries and random gene queries with up to 5 genes from each dataset were generated. The highest TF-IDF scores from the best queries are presented by boxplots and the difference is assessed using a Wilcoxon test.

**Figure S10:**
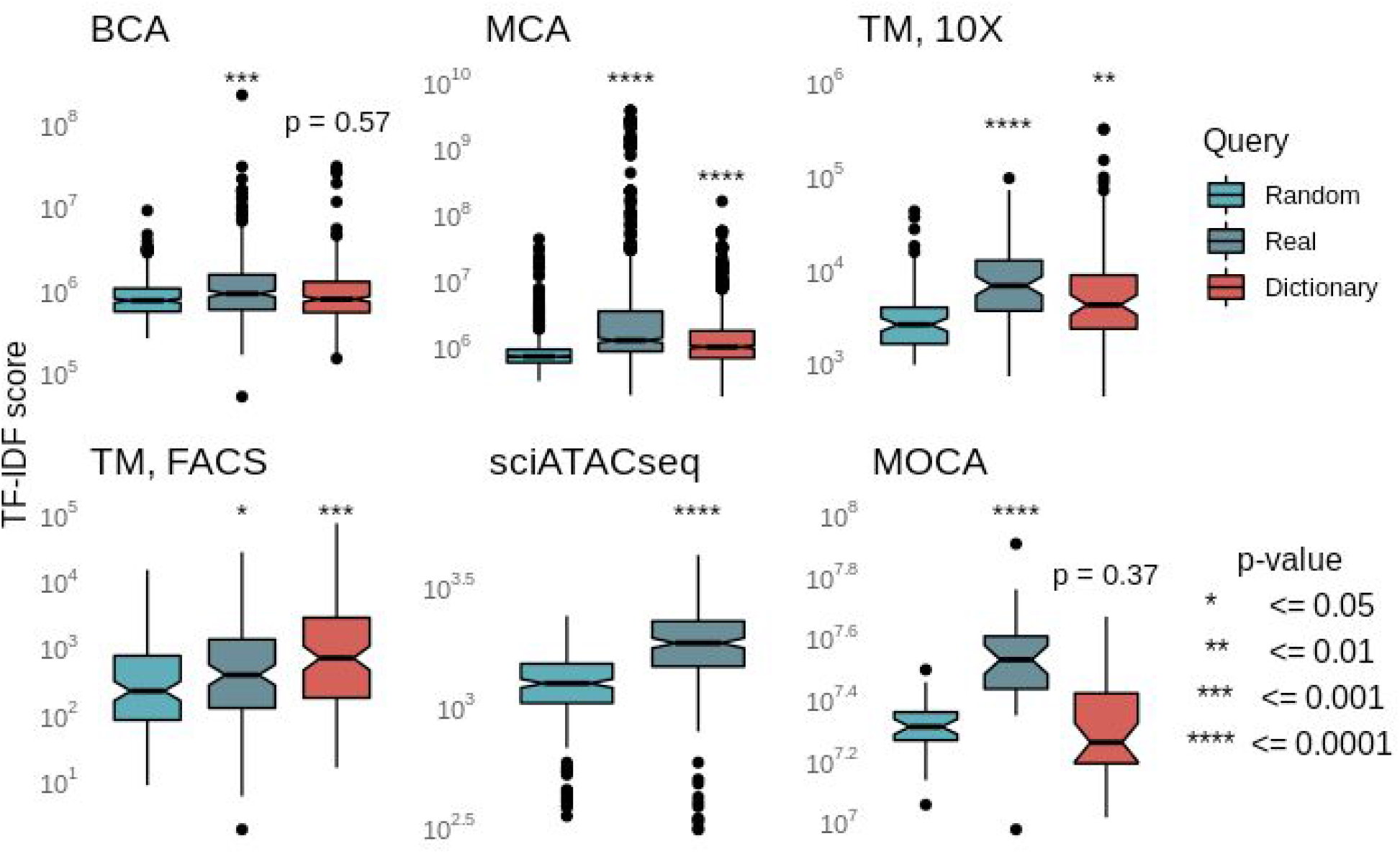
Comparison of TF-IDF score between queries with 10 gene names sampled from each of the top 20 (Real) marker genes, top 1000 (Random) marker genes and phrases to gene names dictionaries (Dictionary) for the Mouse Brain Atlas, Mouse Cell Atlas, Tabula Muris (FACS and 10X), sciATACseq, MOCA. 1000 sets of gene queries per each group were generated. The highest TF-IDF scores from the best queries are presented by boxplots and the difference is assessed using a Wilcoxon test.

**Figure S11:**
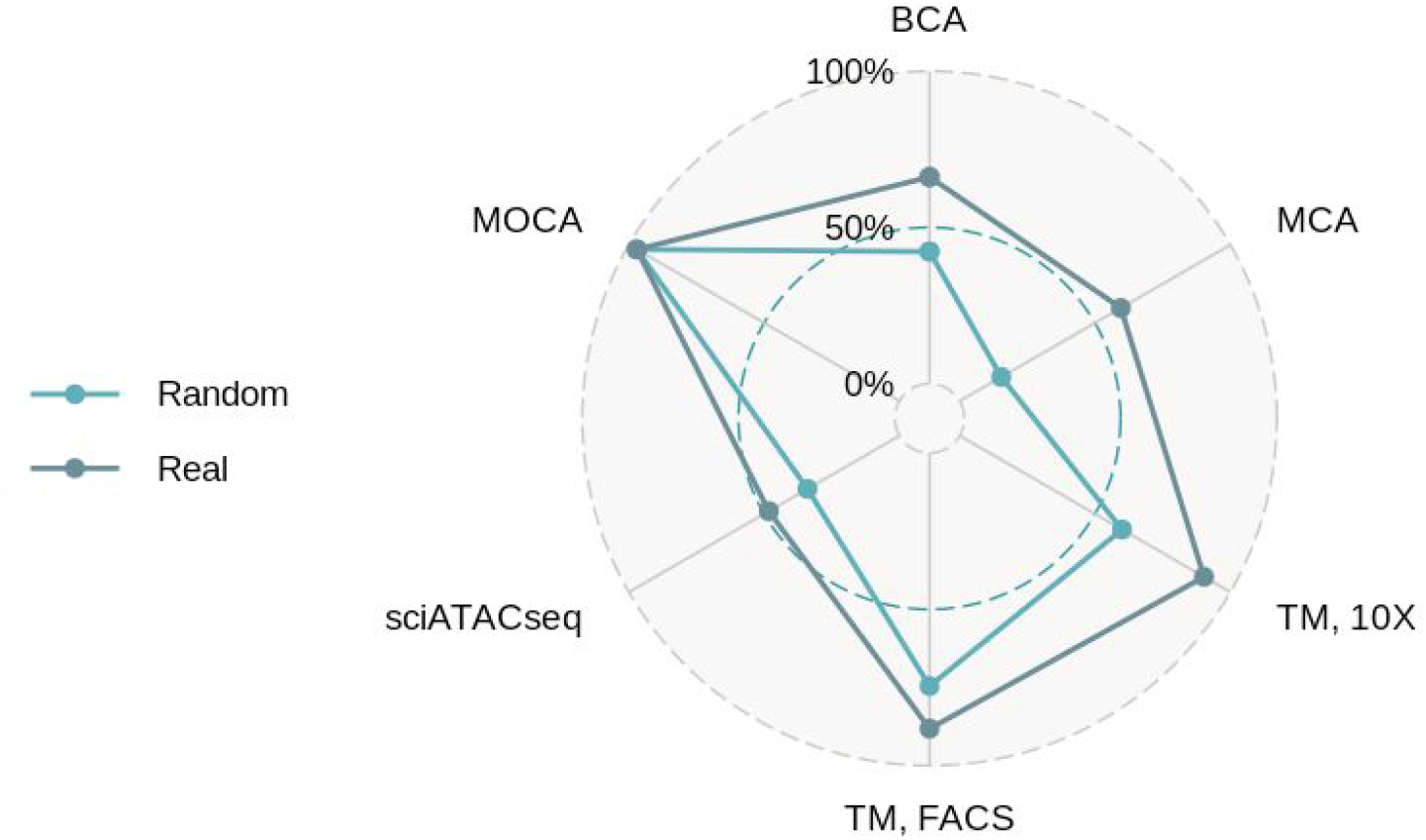
Fraction of searches using the top query from Figure S8 resulting in the desired cell type as the top query.

